# Human voltage-gated Na^+^ and K^+^ channel properties underlie sustained fast AP signaling

**DOI:** 10.1101/2022.11.29.518317

**Authors:** René Wilbers, Verjinia D. Metodieva, Sarah Duverdin, Djai B. Heyer, Anna A. Galakhova, Eline J. Mertens, Tamara D. Versluis, Johannes C. Baayen, Sander Idema, David P. Noske, Niels Verburg, Ronald B. Willemse, Philip C. de Witt Hamer, Maarten H. P. Kole, Christiaan P.J. de Kock, Huibert D. Mansvelder, Natalia A. Goriounova

## Abstract

Human cortical pyramidal neurons are large, have extensive dendritic trees, and yet have surprisingly fast input-output properties: rapid subthreshold synaptic membrane potential changes are reliably encoded in timing of action potentials (APs). Here, we tested whether biophysical properties of voltage-gated sodium (Na^+^) and potassium (K^+^) currents in human pyramidal neurons can explain their fast input-output properties. Human Na^+^ and K^+^ currents exhibited more depolarized voltage-dependence, slower inactivation and faster recovery from inactivation compared with their mouse counterparts. Computational modeling showed that despite lower Na^+^ channel densities in human neurons, the biophysical properties of Na^+^ channels resulted in higher channel availability and contributed to fast AP kinetics stability. Finally, human Na^+^ channel properties also resulted in a larger dynamic range for encoding of subthreshold membrane potential changes. Thus, biophysical adaptations of voltage-gated Na^+^ and K^+^ channels enable fast input-output properties of large human pyramidal neurons.

**One-Sentence Summary:** Biophysical properties of Na^+^ and K^+^ ion channels enable human neurons to reliably encode fast inputs into output.

## Introduction

Cortical computation relies on neurons to process synaptic inputs and transfer these inputs to action potential (AP) output. Electro-morphological properties of neurons shape their input-output function, determine the speed of transfer and how much of the received synaptic information can be passed on as output to downstream neurons. Human excitatory pyramidal neurons in cortical layers (L) 2 and 3 have large dendritic trees receiving over 30,000 synaptic inputs, twice as many as L2/3 pyramidal neurons in rodent cortex (*1*–*4*). Encoding activity of such large numbers of incoming synaptic inputs would require precise temporal resolution through fast input-output processing properties. This fast encoding can be achieved by precise timing of AP output to incoming synaptic input. (Sub-)millisecond precision of AP (spike) timing carries behaviorally relevant information (*5*–*7*), and the ability of neurons to respond to inputs with precisely timed spikes is critical to preserve information and distribute it across synaptically coupled neurons (*8*). While the ability of rodent pyramidal neurons to time APs to synaptic input is limited to 200 Hz (*9, 10*), adult human pyramidal neurons reliably time their APs to subthreshold membrane potential changes of 1000 Hz and above (*11*). Thereby, human L2 and L3 pyramidal neurons have efficient input-output processing properties, with sub-millisecond temporal resolution (*11*). However, mechanisms that enable human pyramidal neurons to reliably encode subthreshold membrane potential changes in AP timing with high temporal precision are poorly understood.

Theoretical studies predict that the speed of AP generation is a critical requirement for fast input-output processing in neuronal networks (*8, 10*). Fast AP onset kinetics allow neurons to respond readily and reliably to fast changing inputs (*12*), while artificial slow-down of the AP waveform impairs the ability of neurons to encode fast inputs, and slows network processing (*10*). In vivo recordings of single neuron activity in human subjects performing cognitive tasks show that neurons increase their firing to 10-50 Hz from almost silent baseline and maintain high firing rates for minutes (*13*–*15*). Typically, in such conditions AP kinetics slow down (*16*) and this could impair precise AP timing and reduce efficient encoding. Neurons with slow AP onset mechanisms will not be able to respond to repetitive synaptic input with an AP, they will ‘ignore’ fast synaptic inputs. Thus, maintaining fast AP shape could be critical for cognitive function.

We recently showed that human neurons are indeed able to maintain fast and stable AP kinetics during sustained firing and this correlates with cognitive function (*17, 18*). Furthermore, the encoding ability of single neurons in awake humans has been linked to cognitive task performance (*19, 20*). Therefore, the ability of human neurons to generate APs with stable shape during repeated firing as well as to encode fast-changing synaptic inputs are inter-related and may play an important role in human cognition. Here we investigated which biophysical mechanisms underlie fast action potential kinetics and input-output encoding in human neurons. To this end, we recorded APs and underlying voltage-gated sodium (Na^+^) and potassium (K^+^) currents from pyramidal neurons in supragranular layers (L2 and L3) of human and mouse association cortices. We characterized biophysical properties of voltage-gated Na^+^ and K^+^ currents underlying action potential generation and used computational modeling to investigate which properties of Na^+^ and K^+^ channels contribute to human AP kinetics and input-output function.

## Results

### Stable and fast human AP waveform during repeated firing

During cognitive tasks such as reading, human neurons in the temporal lobe sustain their AP firing for tens of seconds to minutes at frequencies in the range of 10-50 Hz (*13*–*15*). We first asked whether shape and kinetics of APs in human pyramidal neurons are stable during sustained AP firing, and how this compares to mouse pyramidal neurons. We made whole cell patch-clamp recordings from human and mouse supragranular pyramidal neurons in association cortices and held their membrane potentials to -70 mV. Then we evoked trains of APs with short (3 ms) current pulses at frequencies ranging from 10-70 Hz (Fig. 1A). We observed firing frequency- and time dependent AP broadening, however AP broadening was much more prominent in mouse than human neurons (Fig. 1B-D). Human neurons showed more stable rise and fall speed during repeated firing at all frequencies tested (Fig. 1C-F). Already at a moderate firing frequency of 10 Hz, both rise and fall speed of the 2^nd^ AP showed significantly more slowing down in mouse compared to human neurons (mean relative rise and fall speed, t-test; p=0.013 for rise speed and p=0.016 for fall speed, n=34 human cells, n=32 mouse cells), with more pronounced differences at higher frequencies (Fig. 1F). After 200 APs at 70 Hz the effect was most pronounced: median (Q1-Q3) relative rise speed reduced to 0.56 (0.50-0.67) for mouse neurons (Fig. 1F). In contrast, human neurons sustained their rise speed at a fraction of 0.75 (0.71-0.80) of the initial value, which was significantly higher than in mouse neurons (p=6.2*10^−9^ Wilcoxon rank sum test). The AP fall speed was also significantly more stable in human neurons (mean±SD, human 0.68±0.11, mouse 0.53±0.12, p=3.6*10^−6^, t-test). The initial speed as well as the stability of the rise and fall speed was similar between neurons from epilepsy and tumor patients (Fig. S1) indicating that stability of AP shape does not depend on disease history.

**Fig. 1.**
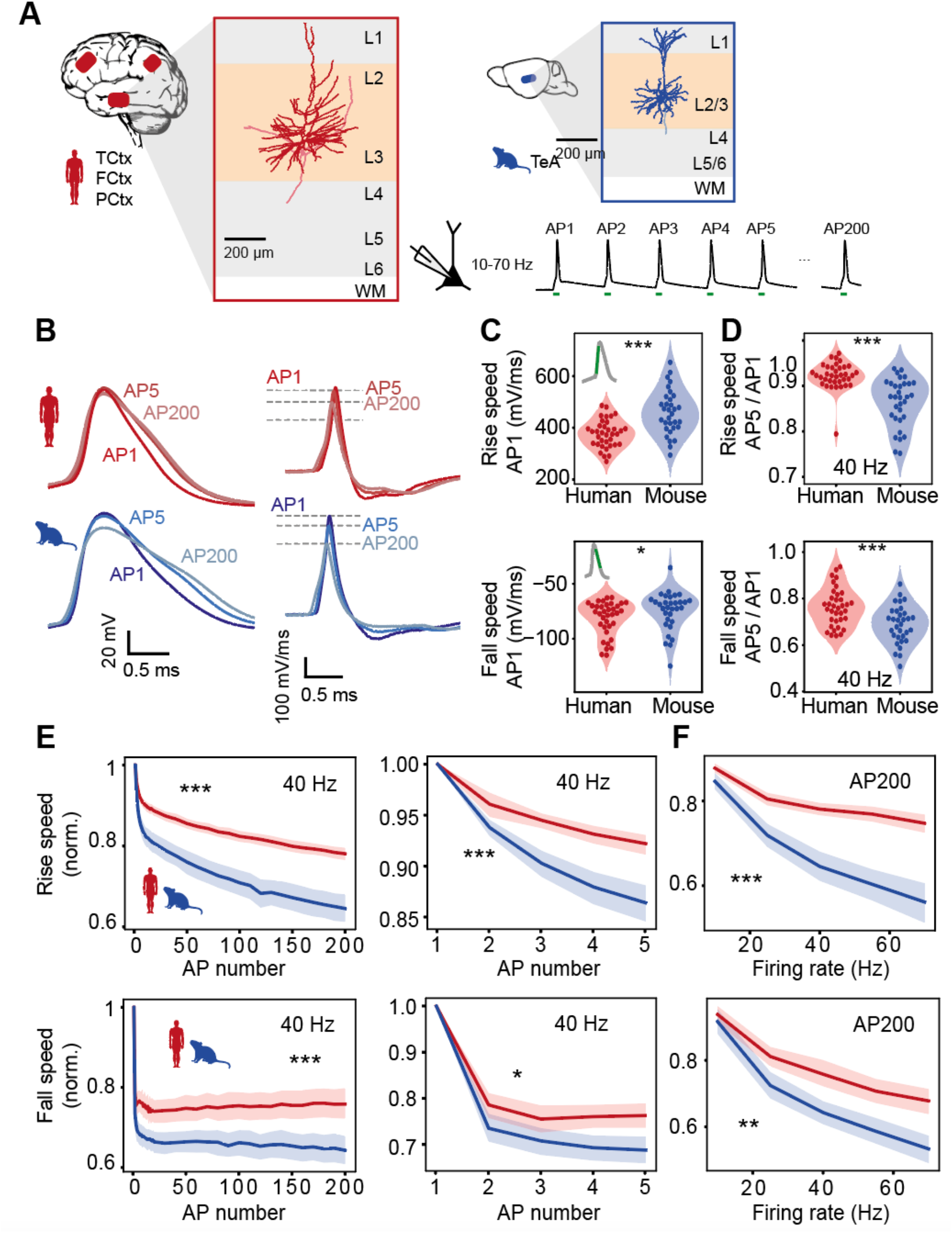
Stable and fast human AP waveform during repeated firing. **(A)** Examples of human (red) and mouse (blue) L2/L3 pyramidal neurons: APs at frequencies ranging from 10 to 70 Hz were evoked by short pulses. (B) Examples of human and mouse AP (left) and AP derivative (right) traces illustrating differences in AP stability. (C-F) AP shape stability in human and mouse neurons (human: n=36 cells, mouse: n=32 cells) (C) Absolute rise and fall speeds of first AP in train. (Rise speed: ***p<10-4, unpaired t-test; Fall speed: *p=0.03, Wilcoxon rank sum test). (D) Relative rise and fall speeds of AP5 normalized to AP1 were faster in human neurons. (Rise speed: ***p<10-6, Wilcoxon rank sum test; Fall speed: ***p<10-3, unpaired t-test) (E) Rise and fall speeds are more stable in human than mouse neurons during repetitive AP firing. Relative rise speed from AP1 to AP200 (left) and AP1 to AP5 (right) during 40 Hz firing showing more slowing of AP kinetics in mouse neurons (significance of Species:AP number interaction in linear regression models: Rise speed AP1-200: ***p<10-26; Fall speed AP1-200: ***p<10-7; Rise speed AP1-5: ***p<10-4; Fall speed AP1-5: *p=0.03). (F) Frequency dependence of rise and fall speed stability after 200 APs (significance of Species:Frequency interaction effect in linear regression models: ***p<10-9, **p=0.001). Shaded areas here and further represent bootstrapped 95% confidence interval of the mean.

### Somatic Na^+ &^ K^+^ currents during action potentials

Since APs mainly arise from a combination of ionic currents through voltage-gated Na^+^ and K^+^ channels (*21*), we asked whether Na^+^ and K^+^ currents showed adaptations during repeated firing. We pharmacologically isolated voltage-gated Na^+^ and K^+^ currents in separate experiments (Fig. S2) and applied realistic, previously recorded AP-train waveforms as voltage command to somatic nucleated patches. When we applied the waveform recorded at 40 Hz in human neurons, the resulting amplitudes of both Na^+^ and K^+^ currents significantly decreased from 1^st^ to the 200^th^ AP (Fig 2A). However, Na^+^ currents in human patches (n=16) reduced substantially less compared to mouse nucleated patches (n=33), and the difference could be observed already during the first 5 APs (Fig. 2B). Furthermore, K^+^ current amplitudes reduced less with subsequent APs in human neurons (n=14, vs. n=11 mouse neurons), but during the first 5 APs there was no species-specific difference in K^+^ current amplitude (Fig. 2B). These effects were independent of whether the applied AP command waveform originated from mouse or human neurons (Fig. S3), indicating that species differences in intrinsic membrane Na^+^ and K^+^ channel properties rather than the shape of the applied AP waveforms are responsible for differences in AP-evoked Na^+^ and K^+^ current adaptation.

**Fig. 2.**
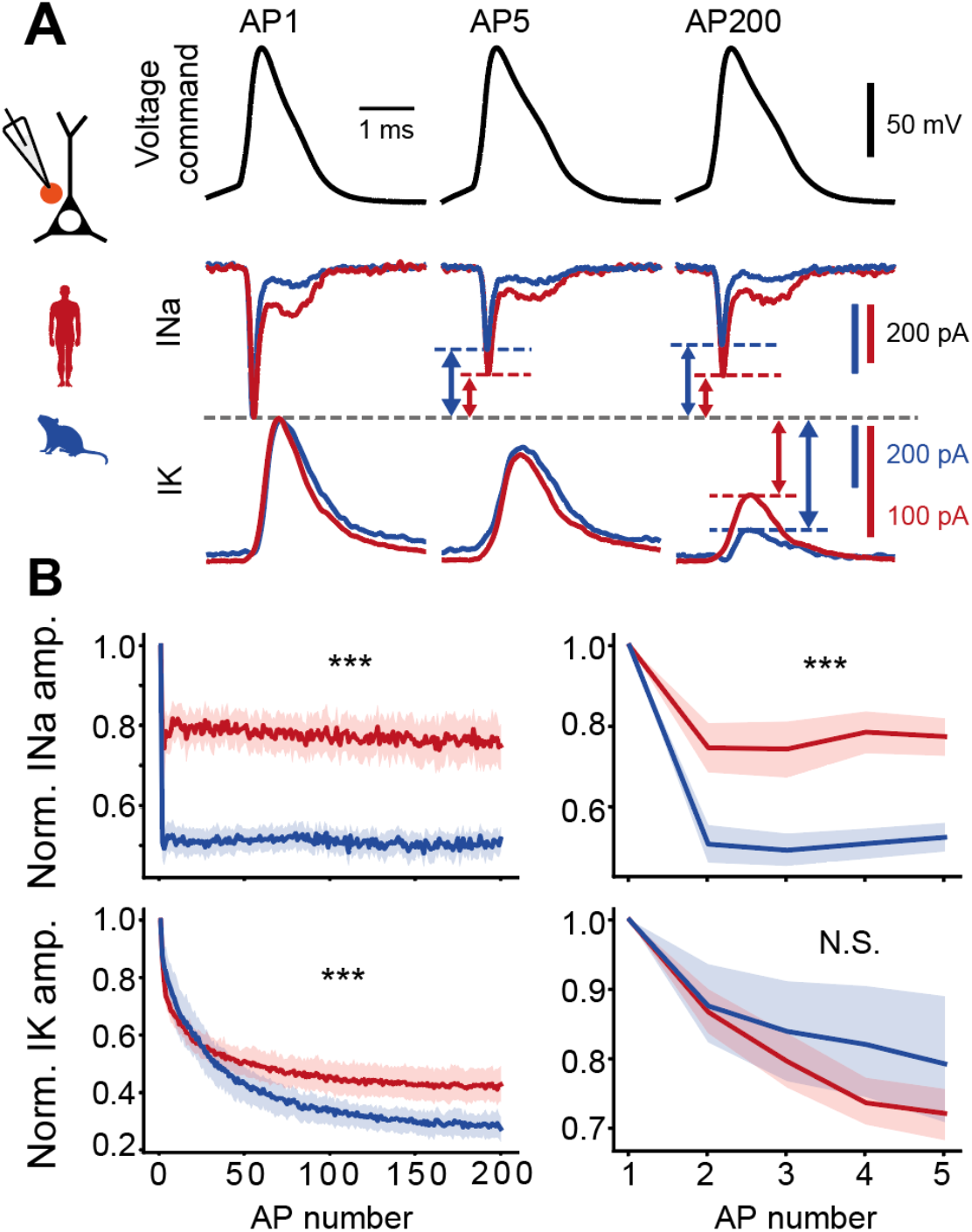
Somatic Na+ and K+ currents during action potentials. (A) Voltage command of human AP waveforms during repeated firing was applied to nucleated somatic outside-out patches and somatic sodium and potassium currents were recorded (separate experiments). Example currents during AP1, AP5 and AP200 at 40 Hz in response to AP waveforms from a human neuron. Arrows and lines depict the relative difference in current amplitude. (B) Mean normalized Na+ and K+ current amplitudes during repeated firing for human and mouse neurons for 200 APs (left) and first 5 APs (right). Asterisks indicate significance of Species:APnumber interaction effect in linear regression model. INa (human: n=16 recordings, mouse: n=33 recordings) amp. 1-200: ***p<10-323, amp. 1-5: ***p<10-53, IK (human: n=14, mouse: n=11) amp. 1-200: ***p<10-45, amp. 1-5: p=0.07.

### Voltage-dependence, activation and inactivation kinetics of voltage-gated Na^+^ channels

To assess which properties of voltage-gated Na^+^ and K^+^ currents explain differences in AP shape stability during AP trains, we tested steady-state voltage-dependence as well as activation and inactivation kinetics of human and mouse voltage-gated Na^+^ and K^+^ currents using square pulses (Figs. 3, 4, S5). We found that both steady-state activation and inactivation curves of Na^+^ currents were right-shifted by respectively 6 and 9 mV towards more depolarized potentials in human neurons (Fig. 3A,B), indicating that at similar voltages a smaller fraction of Na^+^ current was activated and inactivated in human neurons (Table S1, mean±SD V_1/2_ in mV, activation: human= -32.1±4.9, n=19; mouse=-37.9±4.6, n=15, unpaired t-test, p=0.001; inactivation: human= -66.0±7.8, n=18; mouse=-74.8±6.8, n=15, unpaired t-test, p=0.002). At resting membrane potential (estimated at -80 mV due to liquid junction potential, see methods), the average Na^+^ channel availability measured as relative conductance (G/G_max_) was higher (0.81±0.16) for human than mouse neurons (0.63±0.21) (Fig. 3C), while the average resting membrane potential was not different between mouse and human neurons (Fig. S4B). This indicates that compared to mouse, human neurons have a larger fraction of Na^+^ channels ready to be activated during the AP upstroke.

**Fig. 3.**
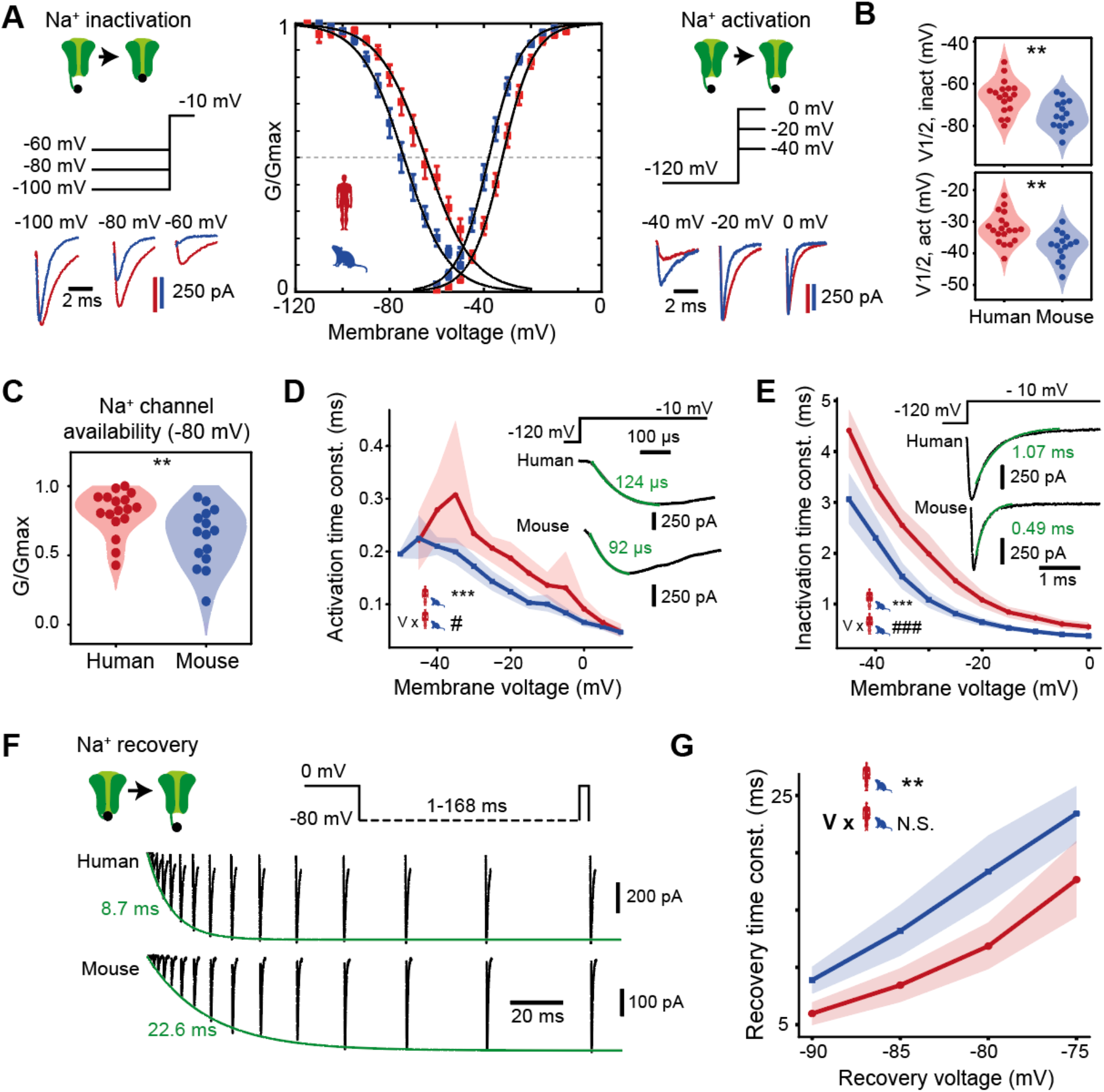
Voltage-dependence, activation and inactivation kinetics of voltage-gated Na+ channels. (A) Na+ currents were evoked following prepulses of incrementing negative voltages (left) and during increasing voltage steps (right) in mouse (blue) and human (red) neurons. Resulting steady-state activation and inactivation conductance curves show a shift to positive voltages for human neurons (middle). Error bars indicate mean ± SEM, black lines are Boltzmann curves fitted to the data. Inactivation: n=18 human recordings and n=15 mouse recordings; Activation: n=19 human recordings and n=15 mouse recordings. (B) Half-activation and half-inactivation voltages are more positive in human neurons (Inactivation: **p=0.002, unpaired t-test; Activation: **p=0.001, unpaired t-test). (C) Relative Na^+^ conductance availability (G/Gmax) at -80 mV is larger in human neurons (n=17 human recordings and 15 mouse recordings). **p=0.006, Wilcoxon rank sum test. (D) Voltage dependence of activation kinetics of Na+ channels in human and mouse neurons: human neurons show longer activation time constants. Inset: example traces of activating Na+ currents during voltage steps to -10 mV, with overlaid exponential fits (green). Sign. of multiple linear regression model: ***p<10-9 for Species effect, #p=0.01 for Species:Voltage interaction effect. (E) Inactivation time constant was longer in human neurons at all membrane voltages. Inset: example currents with fits and time constants shown in green (human: n=19 recordings, mouse: n=15 recordings). ***p<10-21 for Species effect and ###p<10-7 for Voltage:Species interaction effect in polynomial regression model. (F) Example traces of Na+ current recovery at -80 mV at increasing inter-pulse durations; exponential fits to recovery are shown in green. (G) Summary of time constants of Na+ current recovery (shown in F) at different voltages: human neurons showed faster recovery of Na+ currents (human: n=19 recordings, mouse: n=15 recordings). ***p<10-7 for Species effect and N.S. p=0.17 for Voltage:Species interaction effect in linear regression model.

**Fig. 4.**
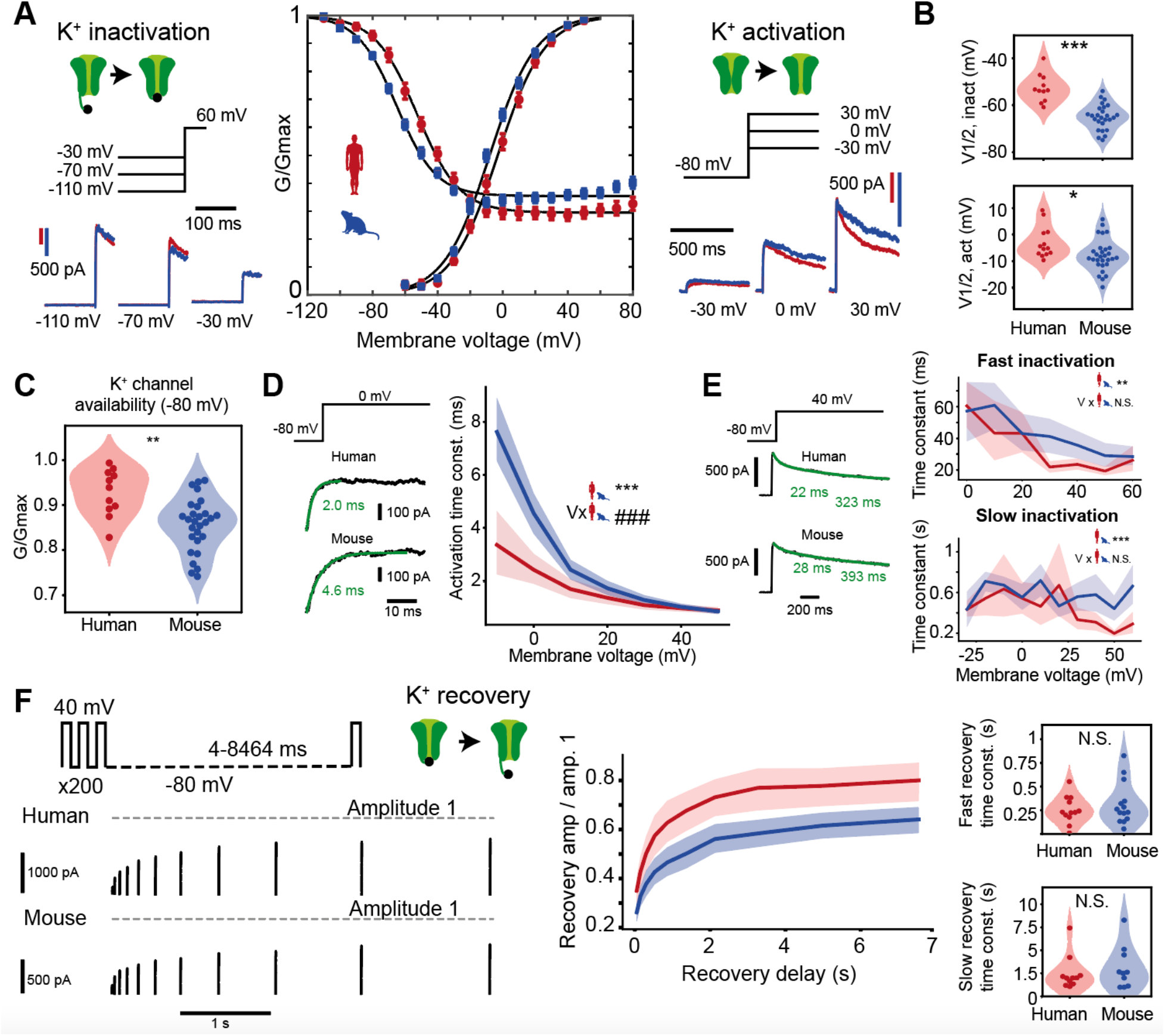
Voltage-dependence, activation and inactivation kinetics of voltage-gated K+ channels. (A) K+ currents were evoked following prepulses of incrementing negative voltages (left) and during increasing voltage steps (right) in mouse (blue) and human (red) neurons. Resulting steady-state activation and inactivation conductance curves show a shift to positive voltage for human neurons (middle). Error bars indicate mean ± SEM, black lines are Boltzmann curves fitted to the data. Inactivation: n=11 human recordings and n=28 mouse recordings; Activation: n=13 human recordings and n=29 mouse recordings. (B) Violin plots of half max-inactivation and activation voltages of individual Boltzmann fits in (A) ***p<10-6, unpaired t-test; *p=0.02, unpaired t-test). (C) Relative K+ conductance availability (G/Gmax) at -80 mV is larger in human neurons (n=11 human recordings and 28 mouse recordings). **p=0.001. Multiple linear regression model: ***p<10-4 for Species effect, ###p<10-3 for Species:Voltage interaction effect. (D) Activation time constant of K+ currents against membrane voltage for both species: K+ currents in human neurons show slower activation at lower voltages. Example traces of activating K+ currents, exponential fits and time constants (green) during voltage steps to 0 mV are shown left. (E) Left: examples of double exponential fits to inactivating K+ current traces, yielding fast and slow time constants. Right: Fast and slow K+ current inactivation time constant versus voltage (n=13 human recordings and 29 mouse recordings). Fast inactivation: **p=0.005, Slow inactivation: ***p=0.0004, Species effect in linear regression models. (F) Example traces of K+ current recovery at - 80 mV at several inter-pulse durations. Red lines indicate response to first pulse. Right, Top: K+ current recovery time courses after recovery at -80 mV with several delays (n=12 human recordings and 14 mouse recordings). Right, Bottom: violin plots for fast and slow recovery time constants. N.S.: not significant, Wilcoxon rank sum test.

The Na^+^ current during the AP upstroke is also dependent on the rate of channel activation and inactivation. To characterize the activation and inactivation rates, we fitted exponential functions to activating and inactivating phases of sodium currents at different voltage steps (Fig 3D,E). At all voltage steps, Na^+^ currents in human neurons had slower activation and inactivation time constants compared with mouse neurons (Fig. 3D,E). Availability of Na^+^ channels for each subsequent AP in the train during repetitive stimulation depends on the rate of recovery of Na^+^ currents from inactivation. To assess the time course and voltage-dependence of Na^+^ currents recovery, we fully inactivated channels by a long depolarizing pulse and applied pulses after increasing time delays at different voltages (Fig. 3F). Human Na^+^ currents had significantly faster recovery time constants at all voltages tested (Fig. 3F, G). In addition, these sodium channel properties lead to more sodium influx during the AP falling phase (Fig. S7, supplementary text). Thus, voltage-gated Na^+^ currents in human neurons activate and inactivate more slowly but recover more quickly from inactivation than their mouse counterparts, which likely results in less accumulation of inactivated channels after an AP (*22, 23*) and faster return to availability to generate the next AP.

### Voltage-dependence, activation and inactivation kinetics of voltage-gated K^+^ channels

Next, we applied square voltage step protocols to nucleated patches and recorded pharmacologically isolated voltage-gated K^+^ currents. The data showed that steady-state activation and inactivation curves (Fig. 4A) were similarly right-shifted towards more depolarized potentials: 5 mV for activation and 12 mV for inactivation (Fig. 4B, Table S1, mean±SD V_1/2_ in mV, Activation: human=-3.0±5.9, n=13; mouse=-7.9±6.0, n=29, unpaired t-test, p=0.02; Inactivation: human=-52.7±6.0, n=11; mouse=-65.0±5.3, n=26, unpaired t-test, p<10^−6^). The shift in voltage-dependence of inactivation resulted in a larger average availability of K^+^ channels at resting membrane potential of human neurons (human: 0.94±0.05, mouse: 0.86±0.06) (Fig. 4C). Human K^+^ currents activated faster than mouse K^+^ currents at lower membrane potentials (Fig. 4D). Inactivation of K^+^ currents had a fast- and slow-inactivating component, which were similar in human and mouse neurons (Fig. 4E). Maximally inactivating K^+^ currents and then applying recovery pulses at -80 mV with various delays showed that, although recovery time constants were similar between species, human K^+^ currents recovered to a larger proportion of the initial amplitude than their mouse counterparts (Fig 4F). Thus, although inactivation and recovery from inactivation kinetics of K^+^ currents were similar between species, a large right-shift of steady-state inactivation resulted in higher K^+^ channel availability. Moreover, a larger recovery from inactivation is likely to aid repolarization during repeated firing in human neurons.

### Human somatic voltage-gated channel properties support stable AP kinetics

Can the differences of voltage-gated Na^+^ and K^+^ current properties explain the differences in stability of the AP in human and mouse neurons? To test this, we simulated the AP command-voltage clamp experiments with Hodgkin-Huxley (HH) conductance-based models for voltage-gated Na^+^ and K^+^ channels with properties determined from experimentally measured steady-state curves and transition rates of both species (Fig S5, Table S2). To isolate the role of conductances in AP generation, the HH-channel models were implemented in single compartment models containing either human or mouse channels (Fig. 5A). Computational simulations with human ion channels showed more stable voltage-gated Na^+^ currents during the first 5 consecutive APs (Fig. 5A,B). Although the model replicated voltage-gated K^+^ currents less well, it showed that similar to the experiments, there were no significant differences between human and mouse K^+^ currents during the first 5 consecutive APs (Fig. 5C). To examine which specific features of the ion channel cause the differences in Na^+^ current stability, we altered mouse Na^+^ channel models by stepwise replacing Na^+^ channel properties with those of human Na^+^ properties and monitored Na^+^ current stability. Endowing mouse Na^+^ channels with human Na^+^ channel activation-related properties did not result in stable Na^+^ current properties of human neurons. In contrast, endowing mouse Na^+^ channels with inactivation properties of human Na^+^ channels produced stability of Na^+^ currents during repeated AP firing replicating the experimentally recorded properties in human neurons (Fig. 5D). Both steady-state voltage-dependence (G/G_max_) as well as time constants of inactivation and recovery contribute to this effect. This suggests that primarily the slow Na^+^ channel inactivation and fast recovery from inactivation are critical for Na^+^ current stability in human neurons.

**Fig. 5.**
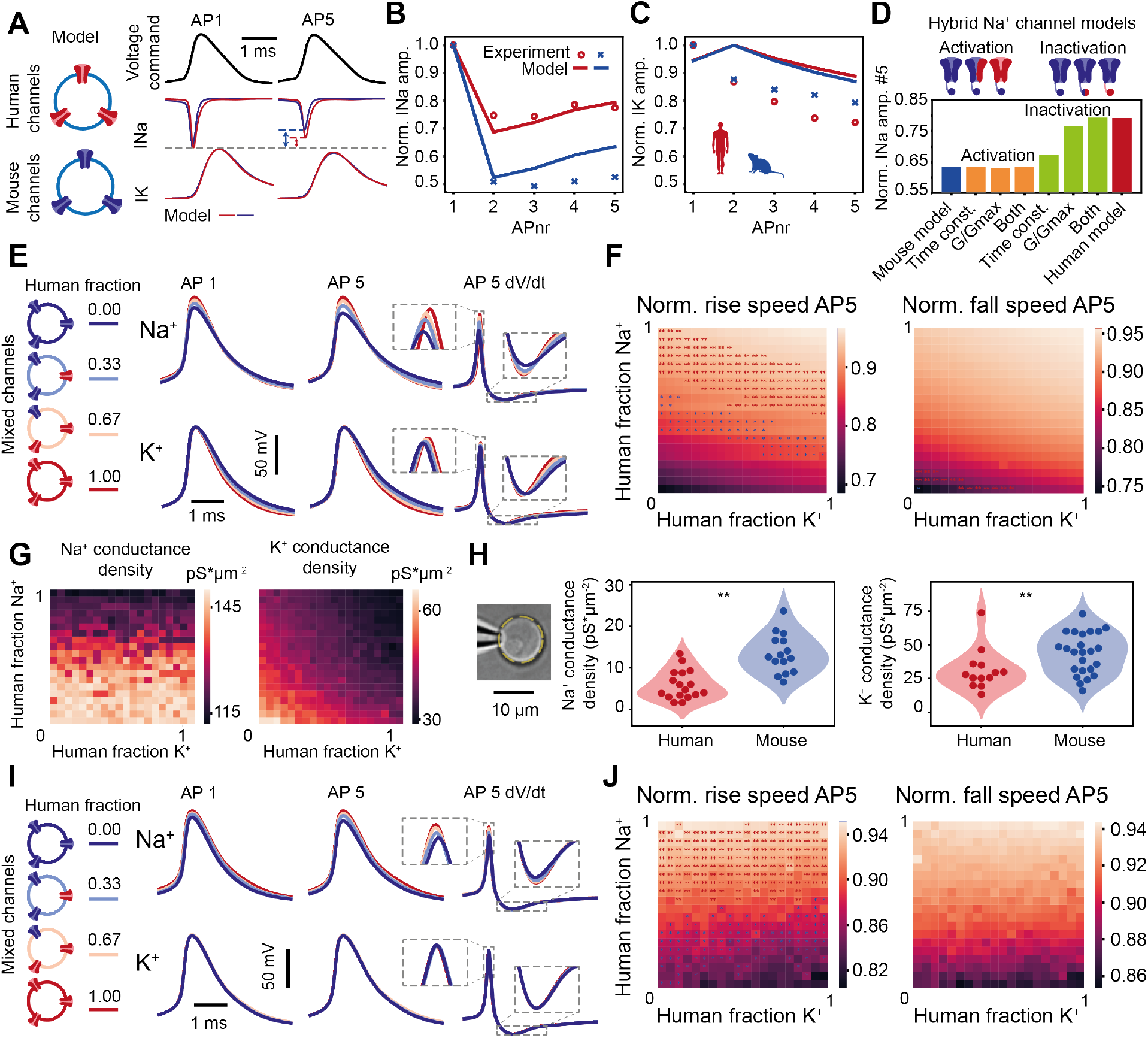
Human somatic voltage-gated channel properties support stable AP kinetics. (A) HH models of human and mouse somatic Na+ and K+ channels based on fits to experimental data. Example traces of simulated Na+ and K+ currents during AP command voltage-clamp experiment. (B,C) Relative amplitudes of simulated and experimentally recorded Na+ and K+ currents. (D) Na+ current stability in different hybrid mouse-human Na+ channel models: stepwise changing different activation and inactivation properties of Na+ channels from mouse to human results in Na+ current stability (current amplitude at fifth AP / current amplitude at first AP). (E) Active AP firing model with variable fractions of human and mouse Na+ and K+ channels. Example traces of first AP, fifth AP and derivative of the fifth AP for various human proportions for Na+ (top) and K+ (bottom) channel simulations. (F) Human/mouse channel fractions affect rise (left) and fall (right) speeds of the fifth AP relative to the first AP. Asterisks indicate results near (±0.02) experimental means from Fig 1D. (G) Na+ and K+ conductance densities at different human/mouse channel fractions. (H) Experimentally recorded Na+ and K+ conductance densities in nucleated patches of human and mouse neurons (Na+ human: n=17 recordings, Na+ mouse: n=15 recordings, K+ human: n=13 recordings, K+ mouse: n=24 recordings) (I), (J) Active AP firing HH-model as in (E,F) but with matched best-fit channel densities from (G).

Next, we asked whether voltage-gated Na^+^ and K^+^ channel properties predicted from the current recordings support AP shape stability. To this end, we simulated AP firing by HH-based single compartment models with varying fractions of human and mouse voltage-gated Na^+^ and K^+^ channels, with constant total channel densities. Both Na^+^ and K^+^ channel properties of human neurons resulted in more stable APs. Gradually replacing mouse Na^+^ and K^+^ channels with their human counterparts increased the speed of both the rising and falling phase of APs (Fig. 5E-F). The amplitude of APs also increased with increasing fractions of human Na^+^ and K^+^ channels relative to mouse channels. Experimentally, however, amplitudes of the first AP did not differ between species (Fig. 1). One explanation for this discrepancy between model and experiment could be that the peak conductance densities are distinct between species. Simulating membrane conductance densities at different fractions of human and mouse voltage-gated Na^+^ and K^+^ channels showed that with larger fractions of human channel properties lower conductance densities were required (Fig. 5G). We next measured experimentally the conductance densities of pharmacologically isolated Na^+^ or K^+^ currents in nucleated patches from human and mouse neurons. Remarkably, both Na^+^ and K^+^ conductance densities were lower in human neurons compared to mouse neurons (Fig. 5H, Table S3, mean ± SEM pS*μm^-2^: human Na^+^ 5.9±0.8; n=17 recordings, mouse Na^+^ 13.4±1.2; n=15 recordings, human K^+^ 31.2±4.2; n=13 recordings, mouse K^+^ 43.5±2.8; n=24 recordings). As reported in the literature, the required sodium conductance density values to generate action potentials in the single compartment computational model exceeded those recorded in the somatic nucleated patch recordings (*24, 25*). Nevertheless, the difference in sodium conductance densities recorded in the somatic nucleated patches (Fig. 5H) support differences in current densities obtained in the model fits (Fig. 5G). Next, we updated the HH models with the obtained fitted channel densities in the models for each specific combination of channel fractions. The simulations revealed that the shape of the first AP was now similar between mouse and human conductance-based models (Fig. 5I), as experimentally measured (Fig. 1C). Rise and fall speeds as well as amplitudes of APs were more stable in models based on increasingly human Na^+^ channel properties. Human K^+^ channel properties did not improve AP stability during the first five APs (Fig. 5I-J). At the 200th AP in the train, human K^+^ channel properties did improve AP stability (Fig. S6). Thus, improved stability of AP shape in human neurons can be explained by properties of their voltage-gated Na^+^ and K^+^ channels, despite lower channel densities.

Human ion channel properties and Na^+^ channel availability support fast input-output conversion Ongoing synaptic transmission results in dynamic sub-threshold dendritic membrane potential changes that may or may not be encoded into AP output (*26*). The extent to which the fine temporal structure of membrane potential changes can be encoded, i.e. how large the bandwidth is of the neuronal transfer function, depends on the stability of AP rise speed properties (*10*). Could the identified biophysical features of human Na^+^ and K^+^ channels underlie the larger encoding properties in human pyramidal neurons (*11*)? To assess the contribution of human and mouse voltage-gated Na^+^ and K^+^ channel properties to the bandwidth of the input-output function of human and mouse neurons, we applied sub-threshold membrane potential changes with varying frequencies to the HH-conductance models with human and mouse Na^+^ and K^+^ channel properties. Model neurons were injected with a DC depolarization to generate APs at frequencies of ∼12-13 Hz, which corresponds to *in vivo* action potential firing by neurons when human subjects perform verbal cognitive tasks (*13, 15, 27*–*29*). On top of the DC component, model neurons received subthreshold sinusoid input currents of increasing frequencies and a Gaussian noise component (Fig. 6A). For each AP, we determined its timing relative to the phase of the injected subthreshold sinusoidal waveform. To quantify phase-locking of AP timing to the sinusoidal input, we generated peri-stimulus time histograms (PSTH) for different sinusoidal input frequencies and calculated modulation depth M/R (peak modulation magnitude M over the mean firing rate R (*11*)) of the best-fit sinusoid to this histogram (Fig. 6B). At similar firing rates in simulations with human and mouse channel properties, M/R was larger, i.e. AP phase-locking was stronger, in models with human channel properties across all frequencies (Fig 6C). When the DC input current level was decreased in the model with mouse channel properties, the firing rate dropped to ∼9 Hz and M/R recovered in the lower frequency range. The cutoff frequency was defined as the input frequency at which the phase-locking (M/R) drops below 0.4. Strikingly, this cutoff frequency was 73 Hz in the model based on human channels, nearly 2-fold higher than 38-42 Hz we obtained in models with mouse channels (Fig 6C). These results suggest that sodium channel properties explain how human neurons can phase-lock their action potential timing to inputs of higher frequencies and broader bandwidth than mouse neurons, as we observed previously (*11*).

**Fig. 6.**
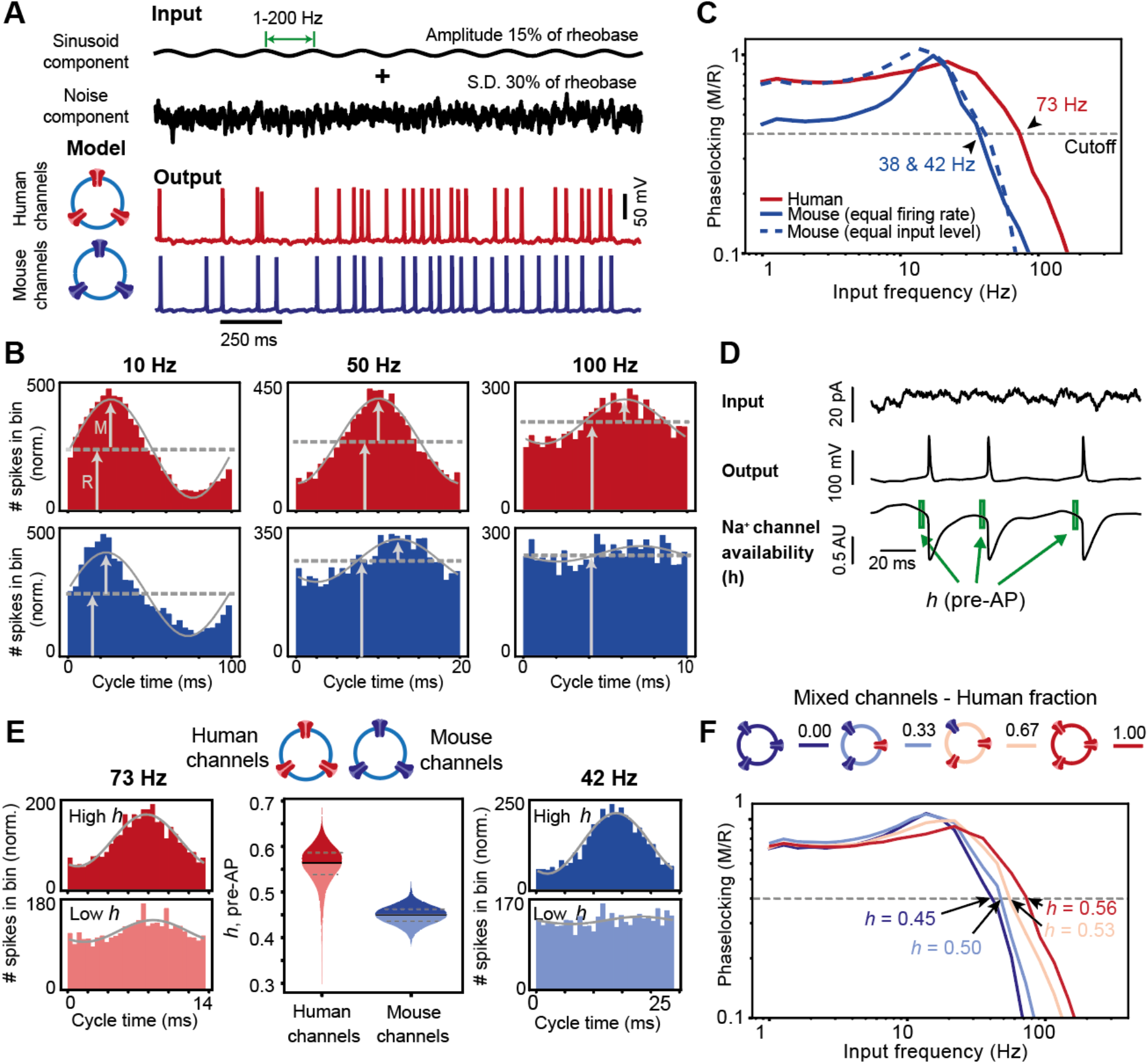
Human ion channel properties and Na+ channel availability support fast input-output conversion. (A) Input current components of AP firing HH-model simulations (top traces, DC component not depicted) and example output traces (bottom traces, input frequency 5 Hz). (B) PSTHs of APs timed within the sinusoid cycle. Grey lines: sinusoidal fit to histogram; black lines: Input sinusoidal current. Arrows indicate mean firing rate R and amplitude M of fitted sinusoid, used to determine the degree of phase-locking (M/R). (C) Phase-locking as function of input frequencies for human and mouse Na+ channel models. Arrows indicate cutoff frequencies. Dashed blue line: mouse Na+ channel simulations with same input current as human Na+ channel simulations (red line). Solid blue line: mouse Na+ channel simulations with input current adjusted to human AP rate of ∼12-13 Hz. (D) Example traces of Na+ channel availability (*h*) resulting from a sinusoid input with added noise. Pre-AP Na+ channel availability is calculated as mean of *h* during 0.8-.2.8 ms before the AP peak (green boxes). (E) Relative timing of APs to sinusoids at cutoff frequencies observed in human (left, input frequency 76 Hz) and mouse (right, input frequency 44 Hz) HH-model simulations with below and above median pre-AP *h* values. Middle panel: Distributions of pre-AP *h* values of all APs in human and mouse channel-based HH-model simulation. (F) HH-model simulations with different fractions of human and mouse voltage-gated channel properties.

Since Na^+^ current properties were largely responsible for stable AP shapes in human neurons (see above), we asked whether species differences in Na^+^ channel inactivation properties would affect the bandwidth of AP phase-locking to input frequencies. To answer this, we quantified for each individual AP the average fraction of sodium channel availability for activation (*h*, see Methods) in the 2 ms preceding AP onset, as well as the AP timing relative to the input sinusoid (Fig. 6D,E). The fraction of available sodium channels (*h*) just before the AP was substantially higher in model simulations with human Na^+^ channel inactivation properties (Fig. 6E, median human *h* = 0.55; median mouse *h* = 0.45). For both human and mouse Na^+^ channel property models, APs that had above median Na^+^ channel availability (*h*) values just before onset were more strictly timed to the input sinusoid than APs with low *h* values (Fig. 6E). Thus, larger sodium channel availability improves phase-locking of AP timing. To test whether sodium channel availability also affected the bandwidth of phase-locking, we varied *h* values in HH-model simulations by varying the relative proportions of voltage-gated channels with human and mouse properties. Increasing the fraction of channels with human properties increased the cutoff frequency of AP phase-locking to sinusoidal inputs to higher frequencies (Fig. 6F), thereby increasing the bandwidth of the input-output transfer function. Thus, properties of human voltage-gated Na^+^ and K^+^ channels support AP shape stability and contribute to improved input-output function of human neurons.

## Discussion

In this study, we identified biophysical mechanisms that can explain fast and stable input-output properties of human pyramidal neurons during repeated AP firing. Both Na^+^ and K^+^ currents had right-shifted steady-state voltage-dependence compared to mouse pyramidal neurons. Voltage-gated K^+^ currents in human neurons activated faster and showed more recovery from inactivation. Voltage-gated Na^+^ currents showed slower and reduced inactivation, as well as faster recovery from inactivation. With model simulations we showed that this resulted in larger functional availability of Na^+^ channels to maintain fast AP onset. Increased availability of Na^+^ channels enable human neurons to time APs to faster subthreshold membrane potential changes, providing these neurons with a larger bandwidth input-output transfer function. These findings provide mechanistic insights into how Na^+^ and K^+^ channels give rise to fast input-output conversion in human pyramidal neurons, the computational building block of the brain.

The rate of AP onset is an important determinant of the ability of neurons to time their APs to inputs (*10, 12, 25*). Properties of voltage-gated sodium channels that favor larger availability of activatable sodium channels are therefore ideally placed to support fast input-output processing. However, voltage-gated sodium and potassium channels are not the only factors. Several recent studies have identified human neuron properties that contribute to fast input-output processing. Firstly, the large dendritic structure of human neurons increases dendritic conductance load and improves AP onset kinetics and phase-locking to high input frequencies (*25*), which was later also confirmed for realistic human morphologies (*17*). Secondly, human L2/L3 pyramidal neurons (*30*) express HCN channels and these channels help speed up the time course at which synaptic inputs arrive at the soma. Lastly, the duration of distal dendritic spikes is dramatically shorter in human neurons when compared to other mammals (*31*).

Although we recorded APs and their frequency-dependent stability in acute slices from neurons containing elaborate axonal and dendritic trees, the voltage-clamp recordings were done in nucleated patches from cell bodies that do not contain the axon initial segment, which is crucial for action potential initiation (*24*). It is likely that differences in voltage-gated channels at the axon initial segment between mouse and human neurons in part also contribute to differences in the AP shape changes we observed in whole-cell recordings. The specific membrane properties and contribution of the human axon initial segment to AP generation remain to be determined. To simulate sodium and potassium currents and their contribution to AP shape changes, we used single compartment models that mimic the somatic nucleated patches we used to record the voltage-gated currents. We previously showed with multicompartment modeling that conductance properties of the somatodendritic compartment can greatly affect the onset rapidity of action potentials initiated at the axon initial segment (*25*). Therefore, our results provide mechanistic insight into how differences in biophysical properties of somatic sodium channels can contribute to differences in AP shape stability between human and mouse neurons.

Sub-millisecond AP timing is not unique in nature. Spike timing with a temporal resolution smaller than the time scales of sensory and motor signals, even at sub-millisecond levels, encode significant amounts of visual information in the fly visual system (*6*). Subcortical neurons in the auditory system can phase lock their firing to auditory frequencies up to ∼1 to 2 kHz (*32*–*34*). However, it is surprising that the large pyramidal neurons in L3 of the human temporal association cortex can time APs with sub-millisecond resolution. It is likely that in evolution several cellular and biophysical adaptations have occurred in human cortical neurons that work in concert to support a fast input-output processing that reliably converts synaptic inputs into timed AP output.

The ability of human neurons to reliably encode highly dynamic subthreshold membrane potential changes with increased reliability in spike timing may allow human neuronal networks to encode information more sparsely (*35*). Although sparse coding has not been compared between humans and rodents, a recent study found that neighboring neurons in human cortex fire less synchronously than in monkey (*36*), suggesting extraordinary sparse coding in human cortex. Indeed, concept neurons in human medial temporal lobe typically respond sparsely, to about 0.5% of concepts (*37*). This would not only increase the number of concepts that can be encoded by a neuronal network (*38*), but would also make the encoding more energy efficient per concept. Thereby there could be a trade-off between energy efficiency at the single neuron level versus energy efficiency at the neuronal network and concept level. We found that reduced Na^+^ channel inactivation and faster recovery from inactivation in human neurons go together with a larger excess influx of sodium during AP firing (Fig S7). While slow or incomplete inactivation of voltage-gated sodium channels during an action potential leads to increased sodium influx and is energetically less efficient (*22, 23, 39, 40*), we here found that these Na^+^ channel properties enable fast input-output processing and improved AP timing to synaptic input. This suggests that while the inactivation dynamics improve information encoding they come at a cost of metabolic efficiency. Recent evidence shows that such a tradeoff exists, since mouse neurons reduce their coding precision to save energy during lack of caloric intake (*41*). At the neuronal network level, coding precision may allow networks to encode with fewer neurons, and thereby overall spend less energy.

Our results show that voltage-gated Na^+^ and K^+^ channels in human neurons have substantially different properties than mouse neurons. Candidate molecular mechanism explaining these differences could be differential expression of Na^+^ and K^+^ channel subunits in mouse versus human neurons. Proteomic studies have shown that in hippocampal synapses several alpha and beta subunits of sodium channels are differentially expressed between rodents and primates (*42*). Genes encoding voltage-gated Na^+^ and K^+^ channels are highly conserved (*43*). However, functionality of the pore-forming alpha subunits can be modulated in several ways, such as auxiliary beta subunits, phosphorylation (*44, 45*), cytoskeleton anchoring and splicing factors (*46*). The molecular basis for the adaptations of Na^+^ and K^+^ channel to support fast input-output processing in human is currently unknown. Although the HH-conductance-based channel models we used to fit human Na^+^ and K^+^ currents capture many biophysical properties of these channels, they did not accurately replicate all properties, such as potassium channel inactivation mechanisms (*47*), which most likely explains the relatively poor fit of the model to the potassium channel currents and action potential fall speeds. Nevertheless, the data and the model simulations combined provide clues pointing to voltage-sensitivity and rate of inactivation properties.

Despite an increased availability of activatable channels, we did observe a reduced density of Na^+^ and K^+^ conductance in human neurons. This is in line with recent work which found that human cortical L5 pyramidal neurons have exceptionally low K^+^ and HCN conductance compared to 9 other mammalian species (*31*). Without biophysical adaptation of sodium and potassium channels, their reduced density would have large consequences for neuronal excitability: an elevated threshold for AP firing and slower input-output processing (*8*). However, our findings on biophysical adaptations can explain why human neurons have fast and stable input-output processing. These include the right-shifted inactivation curves to more depolarized membrane potentials, reduced sodium and potassium channel inactivation and faster recovery from inactivation to maintain a larger proportion of Na^+^ and K^+^ channels available for activation. Our simulations show that these properties can compensate for the reduced conductance density in human neurons, and are vital to understand human cortical function from its neurons up (*48, 49*).

## Materials and Methods

### Human and mouse slice preparation

All procedures related to human patient material were approved by the Medical Ethical Committee of the VU University Medical Center, and in accordance with the declaration of Helsinki and Dutch license procedures. All subjects (N=19; age 21-70, mean±SD age 44.9±17.3 years; 13 males, 5 females, 1 unknown) provided written informed consent for the use of data and tissue for scientific research. All data were anonymized. Human tissue was obtained during neurosurgical removal of healthy cortex which was needed to gain access to a deeper-lying structural pathology: medial temporal sclerosis (N=8) or tumor (N=11). Tissue came from temporal (N=10), frontal (N=5) or parietal (N=4) cortex. Upon removal the tissue was immediately placed into carbogen-saturated NMDG solution (see below) at 0 °C and transported from surgery to the slicing room. There we removed the pia using ultra-fine tweezers and used a vibratome (Leica V1200S) to slice 350 μm thick slices, perpendicular to the cortical surface. Upon slicing in ice-cold carbogenated NMDG solution, each slice was kept for 12 minutes at 34°C in NMDG solution and then transferred to carbogenated holding solution (see below).

All procedures related to mice were approved by the animal ethical care committee of the VU university. C57BL/6 mice (n=43, age 26-74 days, mean±SD age 47.0±13.4 days; 30 males, 11 females, 2 unknown) were anaesthetized with euthasol (i.p., 120 mg/kg in 0.9% NaCl), and transcardially perfused with 10 mL ice-cold carbogen-saturated NMDG solution. Upon removal of the brain, 350 μm thick coronal slices were obtained as described above.

### Solutions

NMDG solution contains (in mM): 93 NMDG, 2.5 KCl, 1.2 NaH_2_PO_4_, 30 NaHCO_3_, 20 HEPES, 25 D-Glucose, 5 Na-ascorbate, 3 Na-pyruvate, 10 MgSO_4_, 0.5 CaCl_2_. pH was adjusted to 7.3 before addition of MgSO_4_ and CaCl_2_ to 7.3 with ∼10-15 mL of 5M HCL. Holding solution contains (in mM): 92 NaCl, 2.5 KCl, 1.2 NaH_2_PO_4_, 30 NaHCO_3_, 20 HEPES, 25 D-Glucose, 5 Na-ascorbate, 3 Na-pyruvate, 2 Thiourea, 2 MgSO_4_, 2 CaCl_2_. pH was adjusted to 7.3 before addition of MgSO_4_ and CaCl_2_ to 7.3 with 1 M NaOH. Recording solution contains (in mM): 126 NaCl, 2.5 KCl, 1.25 NaH_2_PO_4_, 26 NaHCO_3_, 12.5 D-Glucose, 1 MgSO_4_, 2 CaCl_2_. All 3 external solutions were adjusted to 310 mOsm.

K-gluconate internal solution contains (in mM): 110 K-Gluconate, 10 HEPES, 4 KCl, 1 MgATP, 0.3 5’-GTP-Na_2_, 10 Na_2_-Phosphocreatine, 0.2 EGTA, and 20 μg*ml^-1^ glycogen, 0.5 U*μl^-1^ RNAse inhibitor (Takara, 2313A). CsCl internal solution contains (in mM): 130 CsCl, 10 HEPES, 4 MgATP, 0.3 NaGTP, 10 K_2_Phosphocreatine, 2 EGTA. KCl internal solution: as CsCl but all CsCl replaced by KCl. All internal solutions contained 0.5% biocytin.

### Action potential recordings

Pyramidal shaped neurons were identified under DIC within 1200 μm (human) and 400 μm (mouse) from the cortical surface. For action potential recordings (Fig. 1), whole-cell configuration was obtained with 3.5-6 MΩ borosilicate glass pipettes filled with K-gluconate solution, resulting in an access resistance of 7-12 MΩ, which was bridge-balanced in current clamp. Disturbances by synaptic activity were prevented with kynurenic acid (1 mM) and picrotoxin (0.1 mM). Recordings were performed without Bessel filter and digitized at a rate of 125 kHz (Amplifier: Molecular Devices Multiclamp 700B, Digitizer: National Instruments USB-6343). Recordings were obtained using MIES (v1.6, https://github.com/AllenInstitute/MIES) in IgorPro 8 (WaveMetrics) and stored in Neurodata Without Borders format (*50*). Cells were filled with biocytin during recording and after fixation of slices in 4% PFA, cells were stained using a DAB and DAPI protocol to confirm pyramidal morphology and localization in cortical layers 2 or 3. Any recovered cells that did not meet these requirements were excluded from analysis (1 out of 112 recovered cells for the entire study). The rheobase was determined with 10 pA precision by stimulating with 3 ms pulses of increasing amplitude. Then, pulses of 150% rheobase were used to evoke firing at various frequencies. All data presented is without correction for liquid junction potentials (determined experimentally at 15 mV for K-gluconate, and 5 mV for both CsCl and KCl). In these experimental conditions, we interpreted -80 mV (Fig 3C) and -70 mV as resting membrane potential for gluconate and Cl-based internal solutions, respectively.

For all recordings we first determined the rheobase for 3 ms short pulses, that was typically around 1 nA for mouse and 1.5 nA for human neurons. We scaled the short pulses to 150% of the rheobase and this protocol resulted in 100% reliable firing responses (0% failures) even at 70 Hz.

### Action potential analysis

Experimental current clamp traces were filtered off-line using a 20 kHz Butterworth low-pass filter in MATLAB (Mathworks, R2018b). AP threshold was determined as the voltage at the last time the dv/dt crossed 23 mV/ms before the dv/dt maximum. AP rise speeds were directly measured as the maximum of the first temporal derivative of the upstroke of the recorded waveform (*11, 17*) and was quantified as the slope of a linear function fit to the rising phase of the recorded AP voltage trace within a window of 30% to 70% of AP amplitude (defined from AP threshold to AP peak voltage). Both methods had a similar difference between groups (not shown). The AP fall speed was directly measured as the minimum derivative of the downstroke of the recorded waveform (*11, 17*) and was quantified as the slope of a linear function fit to the falling phase of the recorded AP voltage trace within a window of 70% to 30% of AP amplitude. Both methods had a similar outcome (not shown).

### Nucleated patch recordings

For APclamp and channel kinetics experiments, as above but with 2.5-3.5 MΩ pipettes filled with CsCl or KCl internal solution for Na^+^ and K^+^ currents, respectively. Pipette capacitance was greatly reduced by wrapping pipettes in parafilm. After break-in, nucleated patch configuration was achieved by applying 60-80 mbars negative pressure while slowly retracting the pipette. In nucleated patch, access resistance was typically 3-5 MΩ and was compensated in the amplifier using 60-85% prediction and correction. For sodium currents, 1-10 mM 4-AP, 5-10 mM TEA and 1 μM CdCl_2_ were added to external recording solution, for measurements of potassium currents 1 μM TTX and 1 μM CdCl_2_. All recordings were performed at 34 °C, except for sodium channel kinetics, which were done at 25 °C to allow accurate measurement of activation kinetics. For sodium APclamp experiments, P/-5 sweeps were recorded and used to subtract leak and residual capacitive currents. During APclamp experiments for sodium currents the voltage command was 10 mV lower than the original recording in order to account for the 10 mV difference in liquid junction potential between CsCl and K-Gluconate internal solutions. For Na^+^ kinetics experiments, P/N subtraction was done offline using the averaged response to the pre-pulse from -70 to -120 mV.

For experiments in Figure 2 we applied voltage command of previously recorded AP waveform during repeated firing to nucleated somatic outside-out patches and recorded the resulting somatic sodium and potassium currents in separate experiments. We used a voltage command of realistic, experimentally recorded, AP waveforms to mimic the exact voltage changes in the cell that would happen during AP train and elicit sodium currents in response to realistic membrane potential changes.

### Analysis of Na^+^ and K^+^ currents

Na^+ &^ K^+^ currents were analyzed in MATLAB using custom made scripts. Low pass filtering was applied offline to reduce noise, 14 kHz for Na activation, 6 kHz for Na inactivation and 2 kHz for K^+^ traces. To quantify activation and inactivation time constants, exponential functions were fit to the activating and inactivating parts of the recorded current traces. Inactivation time constants were quantified by fitting the data (low-pass filtered to 6 kHz trace during the inactivation phase) to the exponential function: I(t) = a * e^-t/τ^_h_ + b, where I is the current at time t, τ_h_ inactivation time constant, a amplitude and b steady state current (*51, 52*).

Since sodium currents in mouse and human neurons showed large differences in inactivation kinetics, the exponential function to quantify sodium current activation took early sodium channel inactivation into account: I(t) = (1 - e^-(t-d)/τ^_m_) * (a * e^-(t-d)/τ^_h_ + b), where I is the current at time t, d current onset delay, τ_h_ inactivation time constant, τ_m_ activation time constant, a amplitude and b steady state current. A single exponential fit to the activation part of the current, as well as direct quantification of the sodium current activation kinetics using the time to 50% of the maximal amplitude (*53*) yielded similar results (not shown). The ‘goodness-of-fit’ had R-square values above 0.98 for the majority of fits and only fits with R-squared values above 0.9 were included in the analysis.

### Statistical analysis

All statistical analyses were performed in Python, using Scipy and Statsmodels packages. For all data presented in violin plots, normality and equality of variance were tested using d’Agostino-Pearson’s and Bartlett’s tests, respectively. Subsequently, a t-test or Wilcoxon rank sum test was performed to obtain significance levels. For data across voltages, frequencies or APnumbers, confidence intervals of the mean (95%) were bootstrapped and plotted using the Seaborn package. In these cases, statistical analysis was done with linear or polynomial multiple regression models, depending on the best fit.

### Computational models of Na^+^ and K^+^ channels

All computational modeling was performed using the NEURON simulation environment (v7.8) in Python (*54*). Sodium channel models were defined in a similar Hodgkin-Huxley style as in (*55*) (https://senselab.med.yale.edu/ModelDB/ShowModel?model=8210&file=/spikeinit/na3h5.mod#tabs-2), however the steady states were defined by separate Boltzmann equations (as also done for h-gate in (*56*)) to allow assessing of the contribution of steady state and transition rates separately:

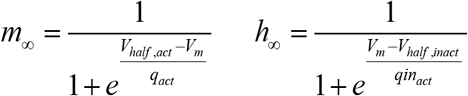

With *m*_*∞*_ and *h*_*∞*_ as the steady states of the activation and inactivation gates, respectively. The model had 14 free parameters: 2 for steady states of each gate, 4 for activation kinetics and 6 for inactivation kinetics). The initial parameters were estimated by fitting the *m*_*∞*_, *h*_*∞*,,_ *m*_*tau*_, *h*_*tau*_ functions to the data. Then the experimental conditions were imitated in NEURON and the exact same voltage clamp protocols were applied, and error was calculated as the sum of squared normalized differences to the experimental means. Subsequently CMA-ES (*57*) was implemented using the DEAP package (*58*) in order to minimize the error over 50-500 evolutionary generations of candidate parameter sets. This process was done separately for the 4 functions and underlying parameters in order to reduce dimensionality problems. The above was repeated independently for both human and mouse data to generate human and mouse channel models.

Potassium channel models were defined and generated similarly as above, with the main exception that the total conductance was calculated as a sum of 3 fractions with different inactivation kinetics, as seen experimentally:

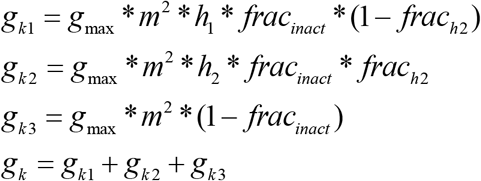

In this scheme, *h*_*1*_ and *h*_*2*_ represent fast and slow inactivation. Since we did not observe voltage modulation of slow inactivation, *h2*_*tau*_ was fixed at experimental values corresponding to inactivation (h_2_ > h_*∞*_) and recovery from inactivation (h_2_ < h_*∞*_). *frac*_*h2*_ was fixed at 0.4, which roughly corresponds to the experimental fraction of slow inactivating current at the highest voltage measured (60 mV). *Frac*_*inact*_ was fixed at 0.85 as this was determined to result in a similar apparent non-inactivating fraction using a 500 ms pre-pulse as observed experimentally. Then the 14 free parameters were fitted on the data same as above for the Na^+^ channel models. The human and mouse K^+^ channel models have the exact same inactivation dynamics, since the experimental time constants of inactivation and recovery from inactivation were not convincingly different enough to justify different models. Therefore, the corresponding experimental means between the species were used to fit the inactivation dynamics. The steady state curves of K^+^ channel inactivation however, were fit separately for each species as they were significantly different between the species.

### APclamp and spiking models

A single-compartment model was generated with diameter of 10 μm, roughly representing a nucleated patch. Membrane capacitance was set to 1 μF*cm^-2^ and axial resistance to 200 Ohm*cm. Reversal potentials of Na^+^ and K^+^ were set to calculated reversals for experimental conditions in current-clamp recordings: 53 and -101 mV, respectively.

The APclamp experiments (Fig. 2) were repeated on the model by implementing a channel model in the NEURON compartment and applying the same voltage waveform as voltage clamp command. This was done with the Na^+^ and K^+^ models of both species as well as several hybrid Na^+^ models. Hybrid Na^+^ models were based on parameter assignments of the mouse Na^+^ model but had the underlying parameters of one or more of the *m*_*∞*_, *h*_*∞*,,_ *m*_*tau*_, *h*_*tau*_ functions assigned to human values.

Then for the spiking models, Na^+^ and K^+^ channels of both species were implemented as well as a passive leak conductance of 1 pS*μm^-2^. In order to gradually change between mouse-like and human-like channel models, global parameters *gmax*_*Na*_, *gmax*_*K*_, *humanfrac*_*Na*_, *humanfrac*_*K*_ were used to set the membrane densities of each channel as below:

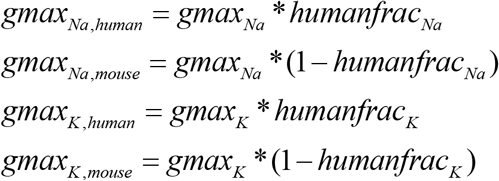

All fits of *gmax*_*Na*_ and *gmax*_*K*_ were fitted using Powell’s method and aimed to minimize the sum of squared differences between modeled amplitude, rise speed and fall speed to cross-species means of experimental values (48.4 mV, 420 mV/ms, 73.1 mV/ms, respectively).

The initial spiking model (Fig. 5E) had all *gmax*_*Na*_ and *gmax*_*K*_ fixed to the best-fit densities for *humanfrac*_*Na*_*=0*.*5* and *humanfrac*_*K*_*=0*.*5*. Then *gmax*_*Na*_ and *gmax*_*k*_ were fitted for each combination of *human frac*_*Na*_ and *humanfrac*_’_ (Fig. 5G), which were then used for spiking models in Fig. 5I.

### Phase-locking

Human and mouse models were initialized as before and with *gmax*_*Na*_ and *gmax*_*K*_ set to their own optimal values (Human, Na^+^: 114.48 pS*μm^-2^ K^+^: 29.05 pS*μm^-2^, Mouse, Na^+^: 148.6 pS*μm^-2^ K^+^: 65.00 pS*μm^-2^). Leak conductance was increased to 2.5 pS*cm^-2^ and reversal potential of sodium was increased to 100 mV, which was necessary to prevent depolarization block during long simulations. Rheobase was determined with 0.1 pA precision and input waves consisting of the sum of a sinusoid and pseudo-random Gaussian noise filtered with a 5-ms exponential filter were generated as described in (*11*). The amplitude of the sinusoid was 15% of the rheobase, and the standard deviation of the noise was 30% of the rheobase. The mean of the sinusoid was used to modulate the firing rate to about 12.5 Hz and was 84% and 94% of the rheobase for the human and mouse model, respectively. For the ‘equal input level’ condition the mean of the mouse sinusoid was lowered to 83%. The seeds to generate the pseudo-random Gaussian noise were kept constant between models to ensure that differences in phase-locking could not be caused by differences in the noise input. For each input frequency, 600 s of time was simulated for each species. Phase-locking (M/R) was calculated as in (*11*), in short the resulting spike times were converted into cycle times by applying the modulus of the cycle duration (*t*_*sprike*_% *F*^−1^), and for each input frequency the PSTH was generated with 30 bins. Then a sinusoid was fitted to the histogram, and the strength of phase-locking was calculated as the amplitude/mean (M/R) of the best-fit sinusoid. The M/R cut-off threshold was set at a value at which the phase-locking was below the low frequency limit at 1-3 Hz in all simulations, which was 0.4. These input frequencies at phase-locking value of 0.4 were used for the simulations in Fig. 6D-F. Using input frequencies at lower cut-off thresholds of 0.3 and 0.2 yielded similar results (not shown).

## Acknowledgements

We would like to thank Dr. Tim Heistek and Ing. Hans Lodder for excellent technical assistance.

## Funding

European Union’s Horizon 2020 Framework Programme for Research and Innovation, specific grant agreement no. 945539 (Human Brain Project SGA3)

ERANET programme iPS&BRAIN

Netherlands Organization for Scientific Research (NWO) Gravitation program BRAINSCAPES: A Roadmap from Neurogenetics to Neurobiology (NWO: 024.004.012)

National Institute of Mental Health (NIMH) award U01MH114812. Netherlands Organization for Scientific Research (NWO) VI.Vidi.213.014 grant

## Author contributions

Conceptualization, R.W., N.A.G., H.D.M. and C.P.J.d.K.

Methodology, R.W., N.A.G. and M.H.P.K.

Software, R.W.

Investigation, R.W., V.M., S.D., D.B.H., A.A.G., E.J.M. and T.D.V.

Formal analysis, R.W.

Funding acquisition, H.D.M.

Resources, S.I., D.P.N., N.V., R.B.W., P.C.d.w.H.

Supervision, N.A.G. R.W., H.D.M. and C.P.J.d.K.

Visualization, R.W. and N.A.G.

Writing – original draft, R.W.

Writing – review & editing, R.W., N.A.G., H.D.M., C.P.J.d.K., M.H.P.K.

## Data and materials availability

All data, model files and scripts used in this work are publicly available at Dataverse repository: https://doi.org/10.34894/L5J0SD.

## Supplementary Materials

Figs. S1 to S7

Tables S1 to S3

