## Supplemental material for "Human voltage-gated Na^+^ and K^+^ channel properties underlie sustained fast AP signaling"

**This PDF file includes:**

Supplementary text
Supplementary Figures S1 to S7
Supplementary Tables S1 to S3

**Supplementary Text**

Increased excess sodium charge during AP firing in human neurons (Fig S7)

$Na^+$ channel inactivation halts sodium influx and prevents excess sodium influx during the falling phase of the AP (*22*). Excess sodium influx is energetically unfavorable since ion gradients need to be restored at the cost of ATP (*22*, *39*, *40*, *59*). The reduced and slower inactivation of human $Na^+$ currents we observed may therefore come at the cost of more energy use. Given that mouse $Na^+$ currents show stronger inactivation and recover slowly from inactivation, we hypothesized that mouse neurons show less excess sodium influx during AP firing (Fig. S7). Indeed, in mouse nucleated patches (n=32), the median excess sodium ratio was 2.3 (1.9-2.7), while in human nucleated patches (n=10) it was 3.8 (3.4-4.1, $p<10^{-5}$, Wilcoxon rank sum test).

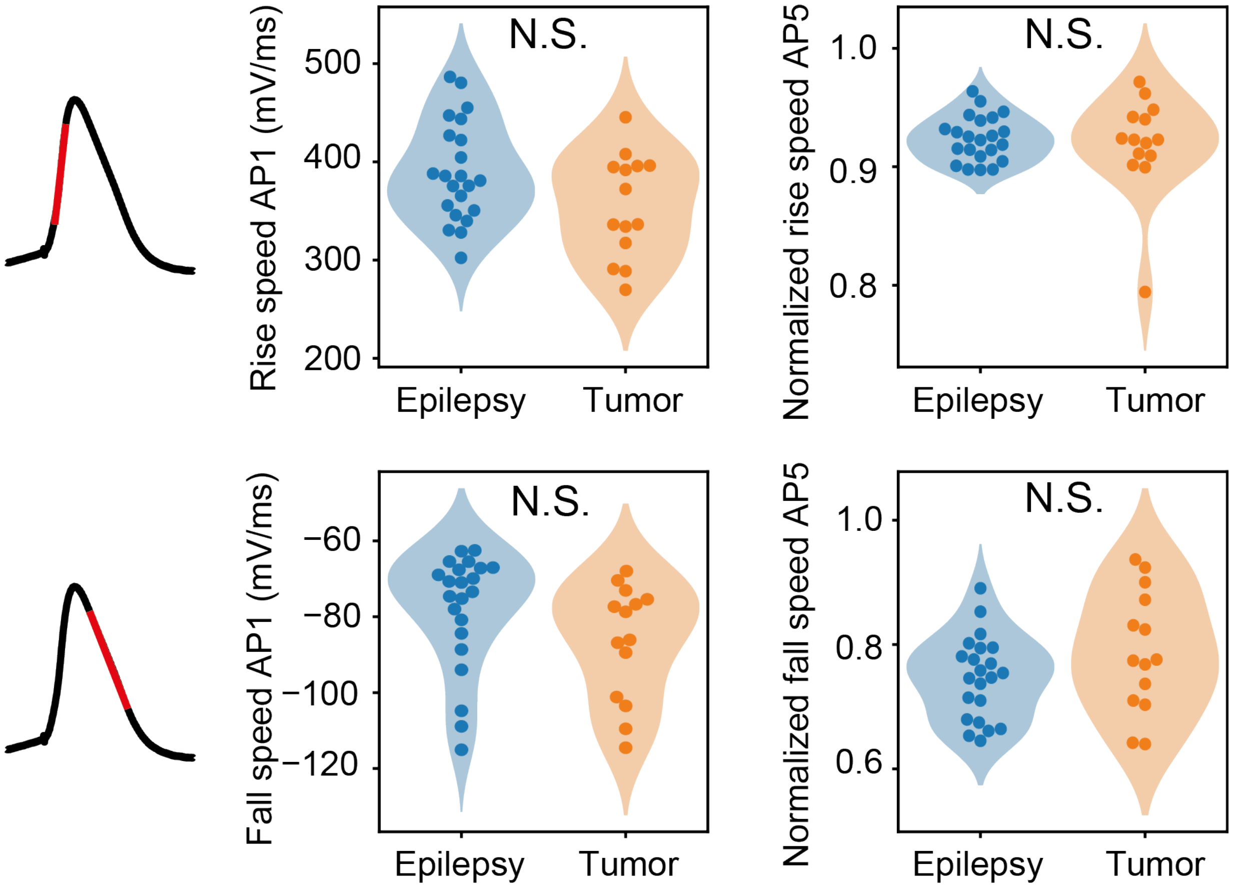


**Fig. S1. Stability of AP rise and fall speed is not related to disease history.** Top: Rise speed of AP1 (left) and normalized rise speed of AP5 (right) for epilepsy patients vs tumor patients. Bottom: AP1 and AP5 fall speed. Statistical evaluation is indicated (t-test or rank-sum test, after multiple comparison correction). N.S.: $P>0.05$

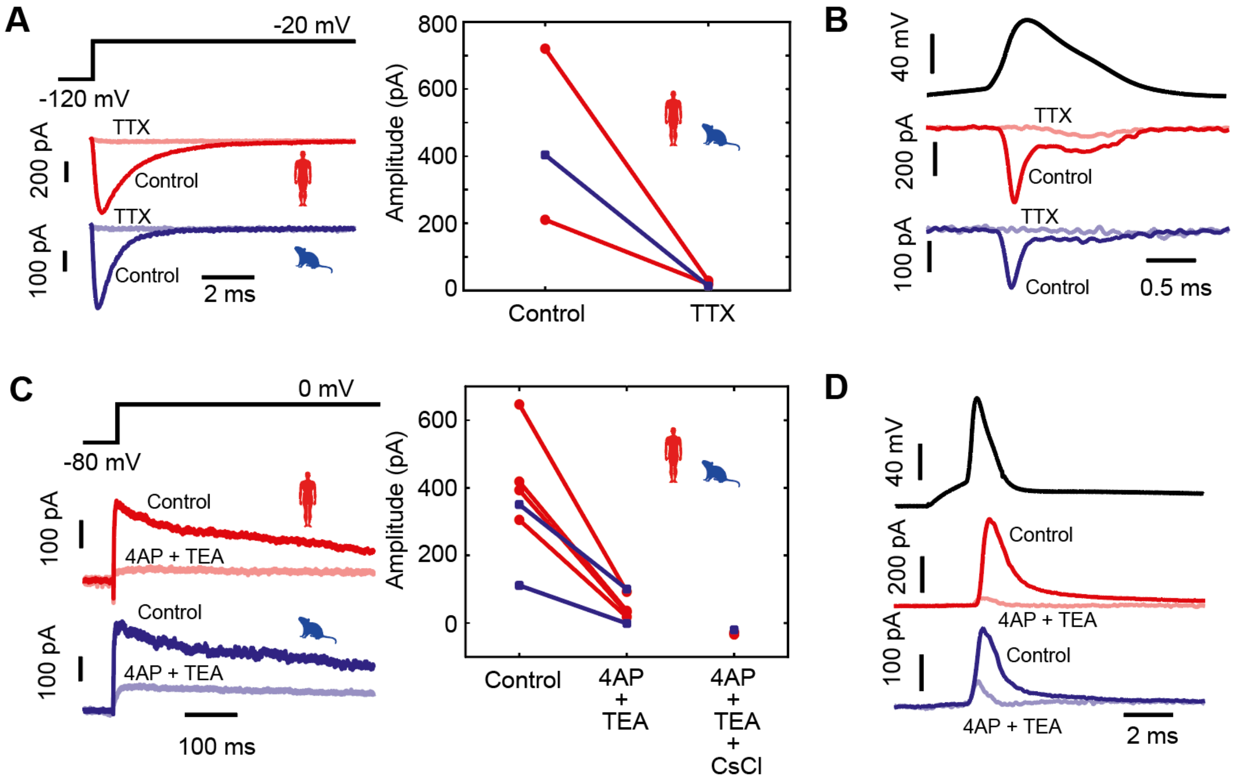


**Fig. S2. Pharmacologically isolated voltage-gated $Na^+$ and $K^+$ currents**. (A) Sodium current example traces before and after application of 1 µM TTX (left) and plotted amplitudes pre vs post application (right). (B) Sodium currents during AP command voltage-clamp experiment before and after application of 10 µM TTX. Distinct set of experiments from (A). (C) Potassium current example traces before and after application of 10 mM 4AP + 20 mM TEA (left) and plotted amplitudes pre vs post application (right). Data from distinct experiment shows complete abolishment of 4AP&TEA -insensitive current using CsCl internal. (D) Potassium currents during AP command voltage-clamp experiment before and after application of 10 mM 4AP + 20 mM TEA.

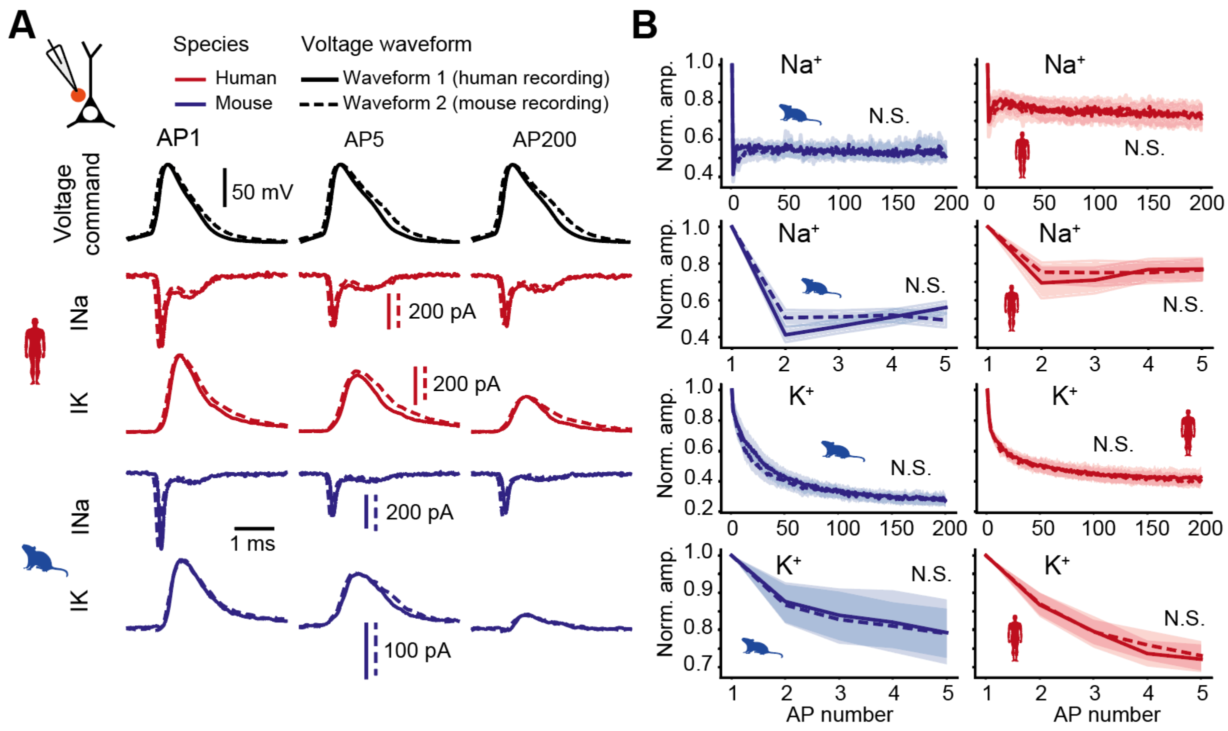


**Fig. S3. Species differences in reduction of repetitive $Na^+$ & $K^+$ currents are not driven by differences in AP command voltage waveform.** (A) Similar example traces of voltage-gated $Na^+$ and $K^+$ currents as in Fig. 2A but with two different AP waveforms. (B) Relative consecutive amplitude of $Na^+$ and $K^+$ currents for both species during different AP waveforms. Plots are shown for both 200 consecutive APs as well as the first 5 APs in separate plots (data from same experiment). Statistical evaluation of AP waveform was done with multiple linear regression models. N.S.: $p>0.05$

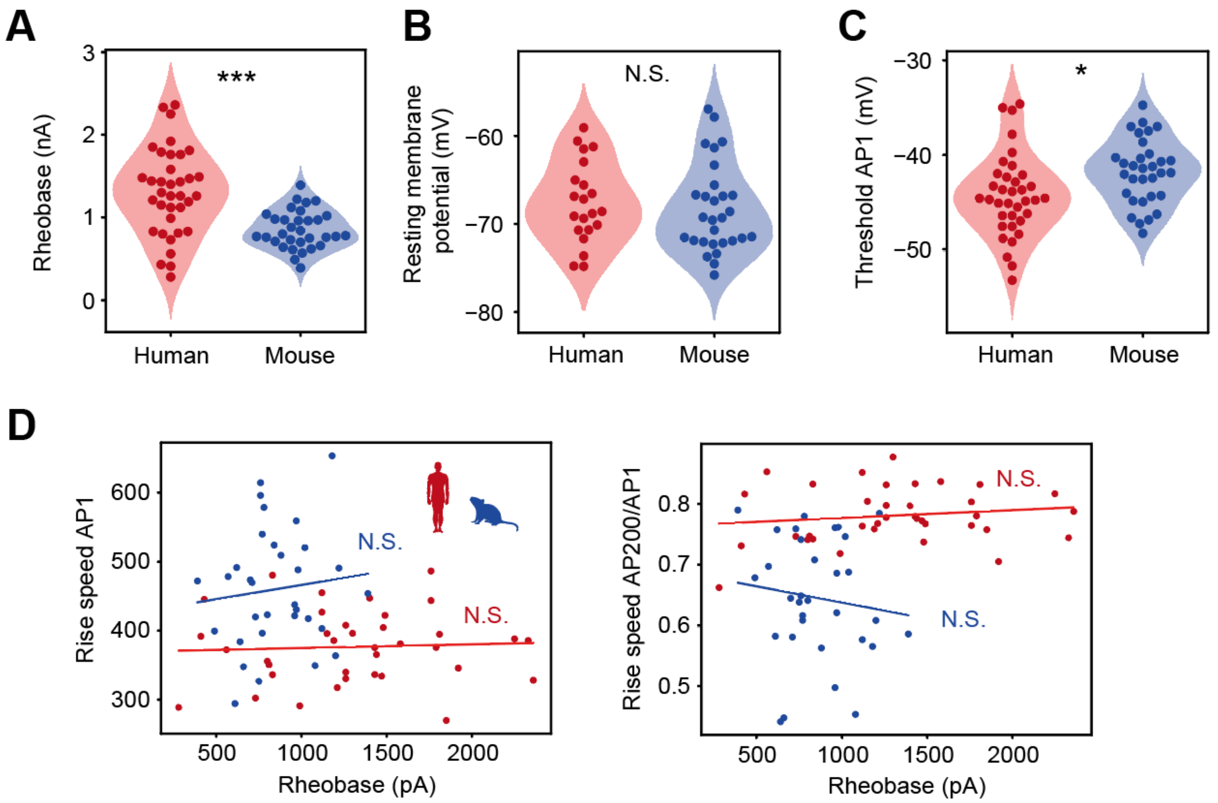


**Fig. S4. AP properties in human and mouse neurons.** (A) Rheobase, minimum current needed to induced AP firing in response to a 3 ms pulse, was higher in human neurons. T-test, p=1.9e-05. (B) Estimated resting membrane potential from neuron's input resistance and injected current to hold the neuron at -70 mV potential. T-test, p=0.59. (C) AP threshold was lower in human neurons. T-test, p=0.011. (D) Rheobase did not correlate with rise speed of AP1 or normalized rise speed of AP200 in mouse or human neurons. Left: human $R^2$=0.003, p=0.76; mouse $R^2$=0.012, p=0.55. Right: human $R^2$=0.021, p=0.4; mouse $R^2$=0.015, p=0.51.

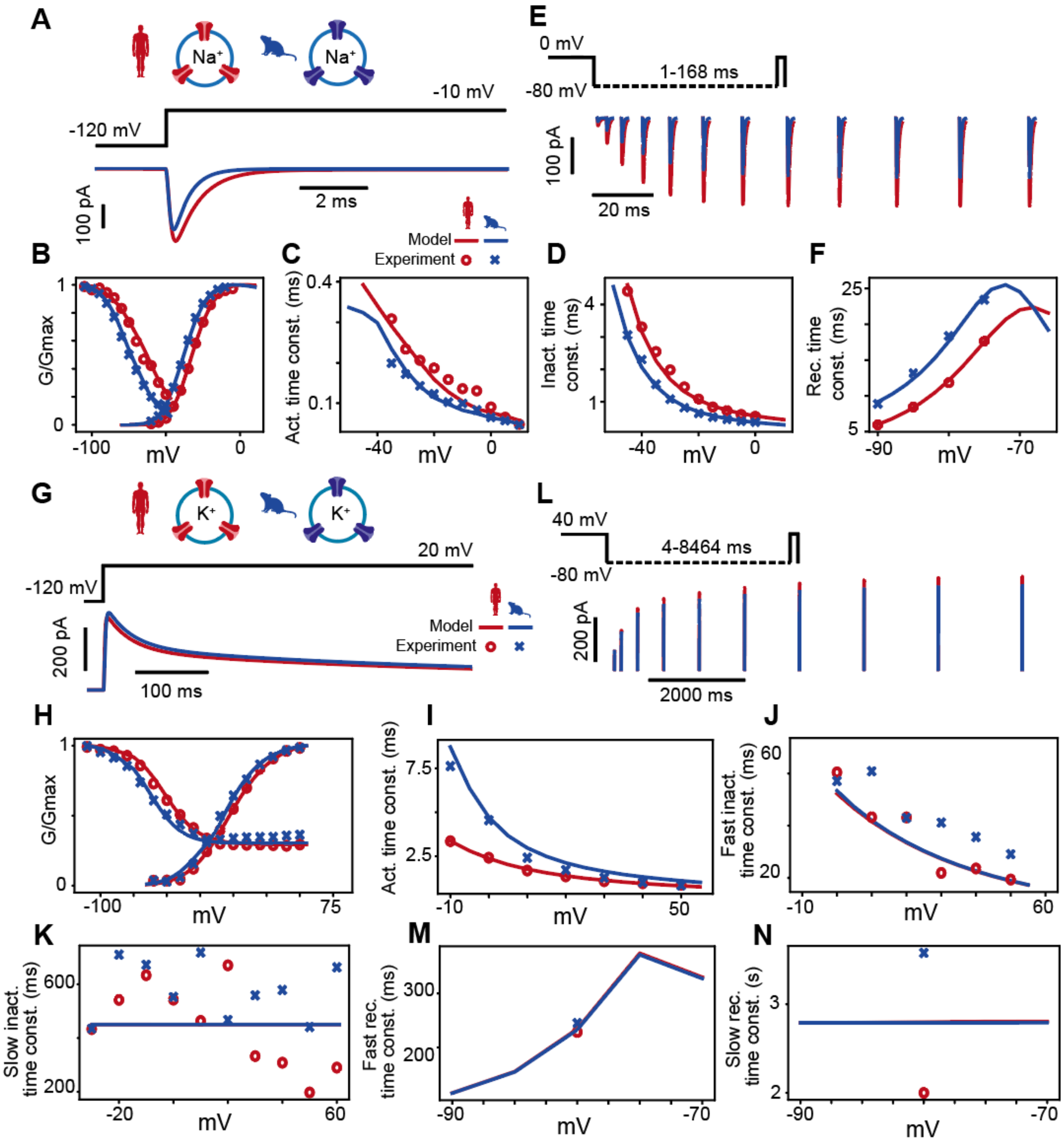


**Fig S5. Models of human and mouse $Na^+$ & $K^+$ channels**. (A) Examples of model $Na^+$ current responses to a -10 mV voltage. (B) Steady state responses of the $Na^+$ models and experimental data are shown. (C) Activation time constants of the experimentally recorded and model $Na^+$ currents. (D) Inactivation time constants of the experimentally recorded and model $Na^+$ currents. (E) Examples of model recovery $Na^+$ currents after variable time delays. (F) Recovery time constants of the experimentally recorded and model $Na^+$ currents. (G) Examples of model $K^+$

current responses to a -10 mV voltage. (H) Steady state responses of the $K^+$ models and experimental data. (I) Activation time constants of the experimentally recorded and model $K^+$ currents. (I) Fast inactivation time constants of the experimentally recorded and model $K^+$ currents. (K) Slow inactivation time constants of the experimentally recorded and model $K^+$ currents. (L) Examples of model recovery $K^+$ currents after variable time delays. (M) Fast recovery time constants of the experimentally recorded and model $K^+$ currents. (N) Slow recovery time constants of the experimentally recorded and model $K^+$ currents. Markers represent means of human (o) and mouse (x) experimental data.

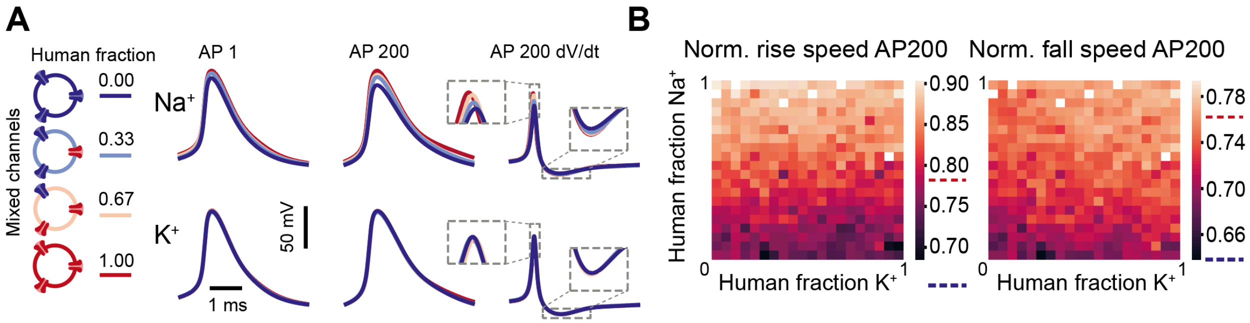


**Fig S6. Human-like $Na^+$ and $K^+$ channel kinetics stabilize APs across 200 APs (A)** Active AP firing model with variable fractions of human and mouse $Na^+$ and $K^+$ channels. Example traces of 1st AP, 200th AP and derivative of the 200th AP for various human proportions for $Na^+$ (top) and $K^+$ (bottom) channel simulations. (B) Quantification of rise (left) and fall (right) speed of AP200 relative to AP1 and dependence on fraction of $Na^+$ and $K^+$ channel models with human properties. Conductance densities as in Fig. 6I-J, with matched best-fit densities from Fig. 6G. Dotted lines next to color legends indicate experimental values for mouse (blue) and human (red) relative rise and fall speed of AP200 at 40 Hz; rise speed: human = 0.78, mouse = 0.65; fall speed: human = 0.76, mouse = 0.64.

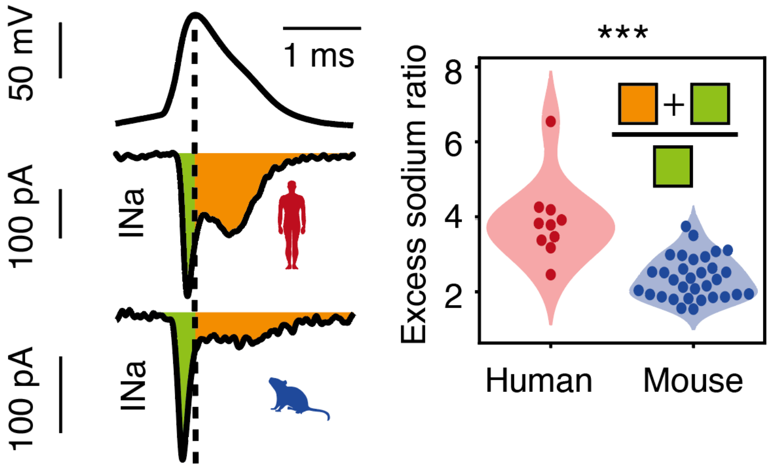


**Fig. S7. Increased excess sodium charge during AP firing in human neurons.** Left: Sodium current traces during AP voltage waveforms in nucleated patches of human and mouse pyramidal neurons. Green and orange shadings indicate the sodium charge before and after the AP peak. Right: Violin plot of excess sodium ratio in both species (human: n=10 recordings, mouse: n=32 recordings). ***$p<10^{-5}$, Wilcoxon rank sum test.

### Supplementary tables

**Table S1. Boltzmann fit characteristics for maximal $Na^+$ and $K^+$ conductances.** Data are not corrected for 15 mV liquid junction potential.

| | **Human** | | **Mouse** | | **Statistics** | | |
|---|---|---|---|---|---|---|---|
| | **mean±SD** | **n** | **mean±SD** | **n** | **Test** | **Statistic** | **p-value** |
| **$Na^+$ activation** | | | | | | | |
| $V_{1/2}$ (mV) | -32.13±4.86 | 19 | -37.88±4.58 | 15 | t-test | 3.513 | 0.0013 |
| Slopefactor ($ms^{-1}$) | 5.77±0.52 | 19 | 5.82±0.43 | 15 | t-test | -0.332 | 0.74 |
| **$Na^+$ inactivation** | | | | | | | |
| $V_{1/2}$ (mV) | -66.00±7.82 | 18 | -74.82±6.81 | 15 | t-test | 3.416 | 0.0018 |
| Slopefactor ($ms^{-1}$) | 7.99±1.92 | 18 | 7.3±0.86 | 15 | t-test | 1.373 | 0.18 |
| **$K^+$ activation** | | | | | | | |
| $V_{1/2}$ (mV) | -3.03±5.95 | 13 | -7.91±6.04 | 29 | t-test | 2.432 | 0.02 |
| Slopefactor ($ms^{-1}$) | 13.87±1.35 | 13 | 13.68±2.17 | 29 | t-test | 0.290 | 0.77 |
| **$K^+$ inactivation** | | | | | | | |
| $V_{1/2}$ (mV) | -52.69±5.97 | 11 | -65.03±5.32 | 26 | t-test | 6.218 | $4*10^{-7}$ |
| Slopefactor ($ms^{-1}$) | 12.8±1.87 | 11 | 11.52±1.97 | 26 | t-test | 1.830 | 0.76 |

**Table S2. Parameters used in NEURON active model.**

| | Human | Mouse | |
|---|---|---|---|
| **General parameters** | | | |
| Diameter (µm) | 10 | 10 | Diameter of the single compartment model |
| Cm (µF*µm$^{-2}$) | 1 | 1 | Specific membrane capacitance |
| Ra (Ω*cm) | 200 | 200 | Axial resistance |
| Rm (Ω*cm$^{-2}$) | 10000 | 10000 | Specific membrane resistance |
| g_pas (S*cm$^{-2}$) | 0.0001 | 0.0001 | Passive conductance density |
| e_pas (mV) | -85 | -85 | Reversal potential of passive conductance |
| gbar_na (pS* µm$^{-2}$) | 114.5 | 148.6 | Maximal $Na^+$ conductance density |
| ena (mV) | 53 | 53 | $Na^+$ reversal potential |
| gbar_k (pS* µm$^{-2}$) | 29.05 | 65 | Maximal $K^+$ conductance density |
| ek (mV) | -101 | -101 | $K^+$ reversal potential |
| **$Na^+$ channel model** | | | |
| th_act_inf (mV) | -42.291457 | -46.6348 | $V_{half}$ for activation steady state |
| q_act_inf (mV) | 10.2311529 | 10.94042 | Slope factor for activation steady state |
| tha (mV) | -58.592356 | -53.6102 | $V_{half}$ for (de)activation time constant |
| qa (mV) | 9.21455372 | 10.08771 | (De)activation time constant slope |
| Ra (ms$^{-1}$) | 0.21398493 | 0.25131 | Open/close rate constant |
| th_act_inf (mV) | -64.48 | -73.97 | $V_{half}$ for inactivation steady state |
| q_inact_inf (mV) | 11.02 | 9.235 | Slope factor for inactivation steady state |
| thi1 (mV) | -39.298677 | 38.18728 | $V_{half}$ for inactivation time constant |
| thi2 (mV) | -77.166745 | 73.39262 | $V_{half}$ for recovery from inactivation time constant |
| qi1 (mV) | 7.21636387 | 7.262844 | Inactivation time constant slope |
| qi2 (mV) | 4.74945005 | 2.771082 | Recovery from inactivation time constant slope |
| Rd (ms$^{-1}$) | 0.04763919 | 0.078592 | Inactivation rate constant |
| Rg (ms$^{-1}$) | 0.01195166 | 0.006373 | Recovery from inactivation rate constant |
| q10 | 2.3 | 2.3 | Temperature sensitivity constant |
| **$K^+$ channel model** | | | |
| th_act_inf (mV) | -18.406164 | -24.9224 | $V_{half}$ for activation steady state |
| q_act_inf (mV) | 19.1995503 | 17.95853 | Slope factor for activation steady state |
| tha (mV) | -29.961798 | -17.1079 | $V_{half}$ for (de)activation time constant |
| qa (mV) | 6.59040569 | 3.688419 | (De)activation time constant slope |
| Ra (ms$^{-1}$) | 0.01845925 | 0.0167 | Open/close rate constant |
| th_act_inf (mV) | -52.61 | -65.2 | $V_{half}$ for inactivation steady state |
| q_inact_inf (mV) | 13.09 | 12.1 | Slope factor for inactivation steady state |
| thi1 (mV) | 40.5659149 | 40.56591 | $V_{half}$ for inactivation time constant |
| thi2 (mV) | -75.938227 | -75.9382 | $V_{half}$ for recovery from inactivation time constant |
| qi1 (mV) | 26.7739292 | 26.77393 | Inactivation time constant slope |
| qi2 (mV) | 0.40183383 | 0.401834 | Recovery from inactivation time constant slope |
| Rd (ms$^{-1}$) | 0.00169014 | 0.00169 | Inactivation rate constant |
| Rg (ms$^{-1}$) | 0.00049688 | 0.000497 | Recovery from inactivation rate constant |
| h2tau_inact (ms) | 450 | 450 | Slow inactivation time constant |
| h2tau_inact_rec (ms) | 2789.79294 | 2789.793 | Slow inactivation recovery time constant |
| prop_inact | 0.85 | 0.85 | Fraction of inactivating conductance |
| prop_h2 | 0.4 | 0.4 | Fraction of slow-inactivating conductance (of prop_inact) |
| q10 | 2.3 | 2.3 | Temperature sensitivity constant |

**Table S3. Surface area and maximal conductance of nucleated patches for $Na^+$ and $K^+$ current recordings.**

| | Human mean±SD | Mouse mean±SD |
|---|---|---|
| **$Na^+$ current recordings** | | |
| Surface area ($\mu m^2$) | 368±63 | 254±32 |
| Conductance $Na^+$ (pS) | 2074±1134 | 3432±1292 |
| $Na^+$ Conductance density (pS* $\mu m^{-2}$) | 5.85±3.38 | 13.39±4.54 |
| **$K^+$ current recordings** | | |
| Surface area ($\mu m^2$) | 307±63 | 228±38 |
| Conductance $K^+$ (pS) | 9398±4399 | 9481±2929 |
| $K^+$ conductance density (pS* $\mu m^{-2}$) | 31.22±14.71 | 43.47±15.04 |
